# Interplay between *ATRX*and *IDH1* mutations governs innate immune responses in diffuse gliomas

**DOI:** 10.1101/2023.04.20.537594

**Authors:** Seethalakshmi Hariharan, Benjamin T. Whitfield, Christopher J. Pirozzi, Matthew S. Waitkus, Michael C. Brown, Michelle L. Bowie, David M. Irvin, Kristen Roso, Rebecca Fuller, Janell Hostettler, Sharvari Dharmaiah, Emiley A. Gibson, Aaron Briley, Avani Mangoli, Casey Fraley, Mariah Shobande, Kevin Stevenson, Gao Zhang, Prit Benny Malgulwar, Hannah Roberts, Martin Roskoski, Ivan Spasojevic, Stephen T. Keir, Yiping He, Maria G. Castro, Jason T. Huse, David M. Ashley

## Abstract

Stimulating the innate immune system has been explored as a therapeutic option for the treatment of gliomas. Inactivating mutations in *ATRX*, defining molecular alterations in *IDH*-mutant astrocytomas, have been implicated in dysfunctional immune signaling. However, little is known about the interplay between ATRX loss and *IDH* mutation on innate immunity. To explore this, we generated *ATRX* knockout glioma models in the presence and absence of the *IDH1^R^*^132^*^H^* mutation. ATRX-deficient glioma cells were sensitive to dsRNA-based innate immune agonism and exhibited impaired lethality and increased T-cell infiltration *in vivo*. However, the presence of *IDH1^R^*^132^*^H^*dampened baseline expression of key innate immune genes and cytokines in a manner restored by genetic and pharmacological IDH1^R132H^ inhibition. IDH1^R132H^ co-expression did not interfere with the *ATRX* KO-mediated sensitivity to dsRNA. Thus, ATRX loss primes cells for recognition of dsRNA, while IDH1^R132H^ reversibly masks this priming. This work reveals innate immunity as a therapeutic vulnerability of astrocytoma.

## Background

Adult-type diffuse gliomas are a diverse group of tumors that account for more than 80% of primary CNS malignancies^1^. They are subclassified based on key molecular alterations, namely isocitrate dehydrogenase (*IDH*) mutation and codeletion of the 1p and 19q chromosomal arms(1p/19q codeletion)^2^, falling into 3 primary groups: 1) *IDH*-wildtype glioblastoma, 2) *IDH*-mutant, 1p/19q co-deleted oligodendroglioma, and 3) *IDH*-mutant, 1p/19q non-codeleted astrocytoma^3, 4^. *IDH*-wildtype glioblastoma is composed primarily of high-grade, clinically aggressive tumors, while both *IDH*-mutant disease groups tend to exhibit lower-grade histopathological features at diagnosis. Treatment for all glioma variants involves some combination of surgery, radiation, and alkylating chemotherapy^5, 6^. However, adult gliomas invariably recur, at which overall prognosis is poor.

Immunotherapy holds considerable potential in the context of central nervous system (CNS) malignancies^7^. Traditional immunotherapies, such as checkpoint blockade, rely on reviving adaptive immune cells like T-cells, and have seen great success in otherwise difficult-to-treat tumors^8^. However, traditional adaptive immune therapies like checkpoint blockade have failed to extend survival in glioma^9^. Despite these disappointments, other unique immunotherapy modalities have shown early promise in the CNS, such as oncolytic viral therapy^10–12^. While oncolytic viruses can selectively lyse tumor cells, their therapeutic potency may be due to their ability to activate the innate immune system through pattern recognition receptors (PRRs)^13, 14^. These receptors detect pathogen associated molecular patterns (PAMPs) and drive multipotent interferon responses, activating multiple arms of the immune system. Prominent examples of PRRs include stimulator of interferon genes (STING), Rig-I-like receptors (RLR), and Toll-like receptors (TLR), with multiple forms of PRR agonism being tested clinically^15–18^.

Inactivation of the SWI/SNF chromatin remodeler gene, α-thalassemia retardation X-linked (*ATRX*), represents a common glioma-associated molecular alteration with the potential to substantially impact the tumor microenvironment^19, 20^. *ATRX* is mutated in more than 80% of *IDH*-mutant, astrocytoma, a large portion of pediatric high-grade glioma (HGG) and a subset of *IDH*-wildtype glioblastoma^19^. Recent studies have demonstrated that loss of epigenetic regulators in general can potentiate responses to immunotherapy^21, 22^. Of note, loss of ATRX appears to alter immune infiltration, cytokine secretion, and chromatin availability to key immune gene regions^23, 24^. Interestingly, *IDH*-mutations, which almost invariably arise with *ATRX* mutations in adult glioma, have been shown to suppress leukocyte chemotaxis and infiltration^25^.

While the association between mutations in *ATRX* and *IDH1* has been known for over a decade, the details of their interaction and basis for frequent co-occurrence in astrocytoma remains unclear. Furthermore, the implication of these mutations on immune-therapeutic responsiveness is understudied. In this study, we leveraged multiple novel glioma models to demonstrate that ATRX deficiency leads to increased innate immune signaling and cytokine secretion in response to innate immune agonism. Furthermore, we found that *IDH1* mutation can mask these pro-inflammatory effects, and that inhibition of mutant *IDH1* relieves this immune suppression. Increases in innate immune signaling and responsiveness to immune agonism led to longer survival in mice with glioma. Taken together, these findings reveal therapeutic potential of targeting the innate immune system for a large, molecularly defined glioma subclass.

## Results

### Astrocytomas are immunologically inflamed relative to oligodendrogliomas

Prior single cell profiling has shown that *IDH* and *ATRX*-mut astrocytomas exhibit increased macrophage and microglial infiltration compared to *IDH*-mut, *ATRX*-WT oligodendrogliomas^26^. To determine if *ATRX* mutations associate with inflammatory signaling pathways in glioma patients, we first performed gene set enrichment analysis (GSEA) on TCGA bulk RNAseq data from *IDH1*-mutant LGGs. The majority of the *ATRX*-mut astrocytomas exhibited enrichment of GO terms associated with immune-related pathways, including activation of innate immune response, PRR activation, and response to IFNα and β (Fig. 1a; Supplementary Fig. 1a-b). By contrast, *ATRX*-WT/ 1p-19q co-deleted oligodendrogliomas exhibited weaker correlations with these immune-related transcriptional signatures. With the notable exception of loci included in the oligodendroglioma-defining 1p/19q codeletion event (*IRF3*, *JAK1*, *ISG15, IFNL1,* and *IFNL2*) we found a general paucity of mutations, high-level amplifications, and/or deep deletions associated with *ATRX* mutations (Supplementary Fig. 2), indicating that the effects of glioma-associated molecular alterations like *IDH* mutation and ATRX deficiency are likely not mediated via genetic mutations or copy number variations.

**Figure 1:**
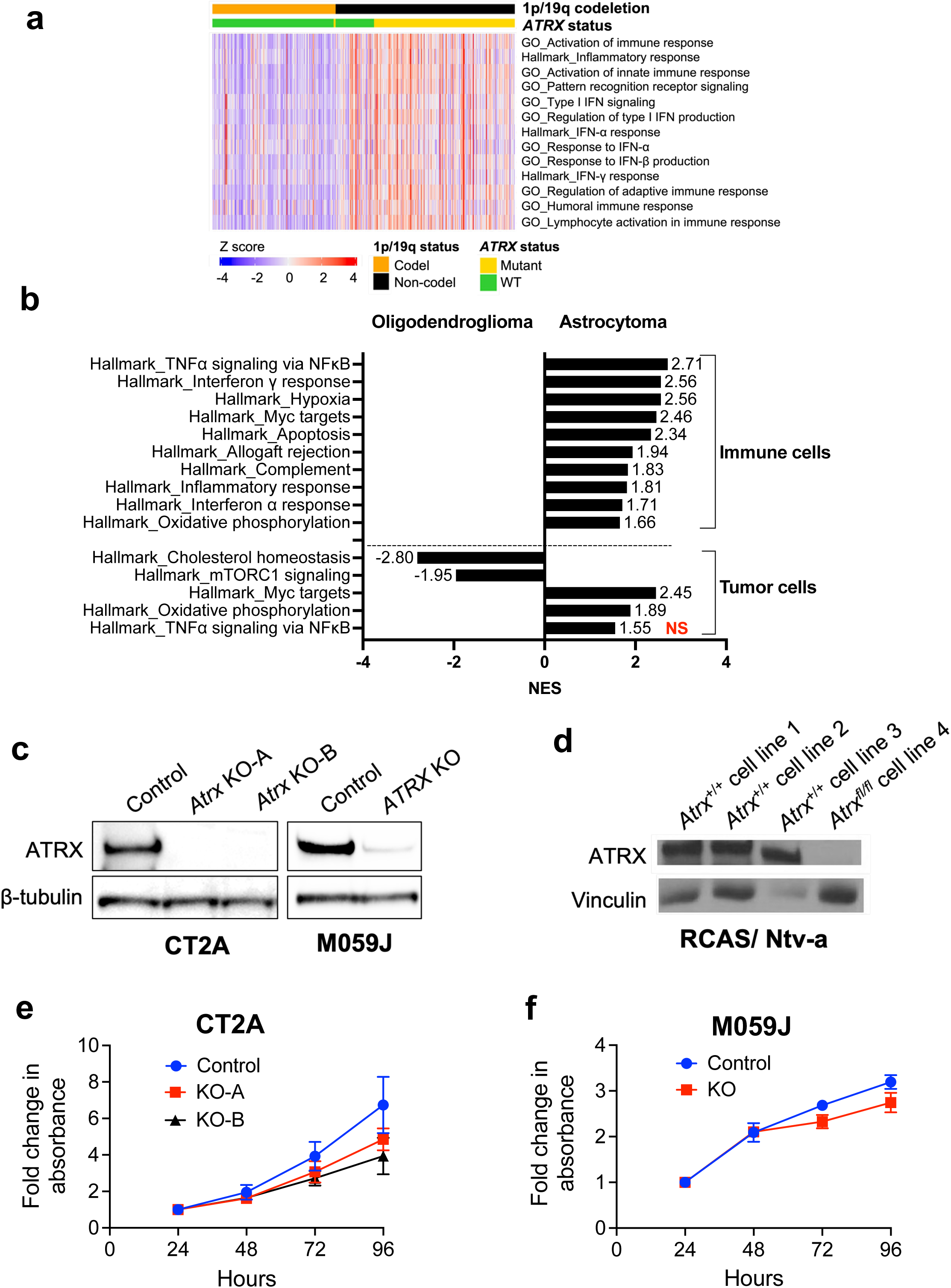
Astrocytomas are immunologically engaged compared to oligodendrogliomas. (**a**) Heatmap from ssGSEA showing enrichment for various innate immune related gene sets in 1p-19q noncodel/ *IDH* mutant/ *ATRX* mutant astrocytomas (n=191) compared to 1p-19q codel/ *IDH* mutant/ *ATRX* WT oligodendrogliomas (n=164). A relatively smaller number of 1p-19q codel/ *IDH* mutant/ *ATRX* mutant (n=3) and 1p-19q noncodel/ *IDH* mutant/ *ATRX* WT (n=52) are also included in the analysis for comparison. (**b**) GSEA Hallmark gene sets of scRNAseq data from immune cell clusters and tumor cell clusters from *IDH* mutant oligodendroglioma (n=6) and *IDH* mutant astrocytoma (n=10) patient tumor samples. Normalized enrichment scores (NES) for gene sets that have adjusted p value <0.05 (except tumor cells – Hallmark_TNFα signaling via NF^κ^B highlighted in red) are shown. (**c**) Western blot using lysates from ATRX-intact and ATRX-depleted mouse CT2A cells and human M059J cells showing total ATRX expression. KO-A and KO-B represent two ATRX-deficient CT2A single cell clones used in this study. β-tubulin serves as the loading control. (**d**) Western blot using lysates from RCAS/Ntv-a cell lines derived from 3 separate tumor bearing *Atrx*^+/+^+ Cre mice (cell lines 1, 2 and 3) and one *Atrx^fl/fl^* + Cre mouse (cell line 4) showing total ATRX expression. Vinculin serves as the loading control. (**e, f**) Growth assays of CT2A CRISPR control (Ctrl) or *Atrx*-KO cells (KO-A, KO-B) (**e**) and M059J CRISPR control (Ctrl) and *ATRX*-KO (KO) cells (**f**) in culture for 96 hours. Data indicates fold change values normalized to corresponding 24hrs absorbance value for each cell line shown as mean ± SEM; n=3 biological replicates for CT2A and n=2 biological replicates for M059J.

To determine the relative contributions of tumor (glioma) vs immune cells in driving the differential inflammatory signatures noted in Fig. 1a, we queried single cell RNAseq data^26, 27^ from *IDH*-mutant oligodendrogliomas and astrocytomas. Relative to Oligodendrogliomas, in *IDH*-mutant astrocytomas, immune cell clusters were enriched for gene sets associated with inflammatory response, IFNα response, IFNγ response and TNFα signaling via NF1B; whereas tumor cell clusters exhibited minimal enrichment for immune related molecular networks (Fig.1b). This lack of enrichment for immune related gene sets may be a result of expression of the immunosuppressive *IDH1^R132H^* in astrocytic tumors, while the tumor microenvironment is still immunologically active. Overall, these data echo earlier findings^26^, pointing to a higher degree of immune cell infiltration in *IDH*-mut, *ATRX*-mut astrocytomas relative to *IDH*-mut, *ATRX*-WT oligodendrogliomas. These findings may indicate a role for ATRX deficiency in promoting an immune-reactive phenotype.

### Generation of ATRX-depleted glioma models

To investigate the role of ATRX deficiency in regulating inflammation in gliomas, we developed both mouse and human experimental systems with intact immune signaling pathways. Using a CRISPR/Cas9 approach, we generated single cell *Atrx*-KO clones—KO-A and KO-B—isolated from the murine CT2A glioma cell line and a polyclonal *ATRX*-KO derivative of the human M059J glioma cell line, along with corresponding controls. Additionally, we developed a genetically engineered murine glioma model, leveraging RCAS/Ntv-a retroviral transduction to express the oncogene platelet-derived growth factor a (PDGFA), an shRNA targeting *Tp53*, and Cre-recombinase in nestin-positive cells within the brains of isogenic mice harboring either intact or floxed *Atrx* loci^28^. This strategy yielded ATRX-intact and deficient gliomas that could then be subjected to *ex vivo* culture as well as serial transplantation. All ATRX-deficient murine and human cell line reagents demonstrated robust depletion of ATRX protein (Fig. 1c, d). Interestingly, ATRX deficiency modestly impaired *in vitro* growth (Fig. 1e, f).

### ATRX loss impairs glioma growth *in vivo* in a manner largely dependent on the immune microenvironment

Next, we evaluated the extent to which ATRX inactivation impacts glioma growth *in vivo*. Abundant genetic evidence supports the classification of *ATRX* as a tumor suppressor in human cancer^29^. However, the majority of adult gliomas harboring ATRX deficiency exhibit relatively indolent biology at initial diagnosis despite inexorable malignant progression over time^30^. Consistent with this behavior, we found that *Atrx* knockout was associated with extended survival in immunocompetent mice subjected to orthotopic allografting with CT2A cell derivatives (Fig 2a; Supplementary Fig. 3a). Moreover, mice harboring ATRX-deficient gliomas in the context of our RCAS/Ntv-a genetically engineered model also exhibited extended survival relative to ATRX-intact counterparts, in both *de novo* and re-injection contexts (Fig 2b; Supplementary Fig. 3b-c).

**Figure 2:**
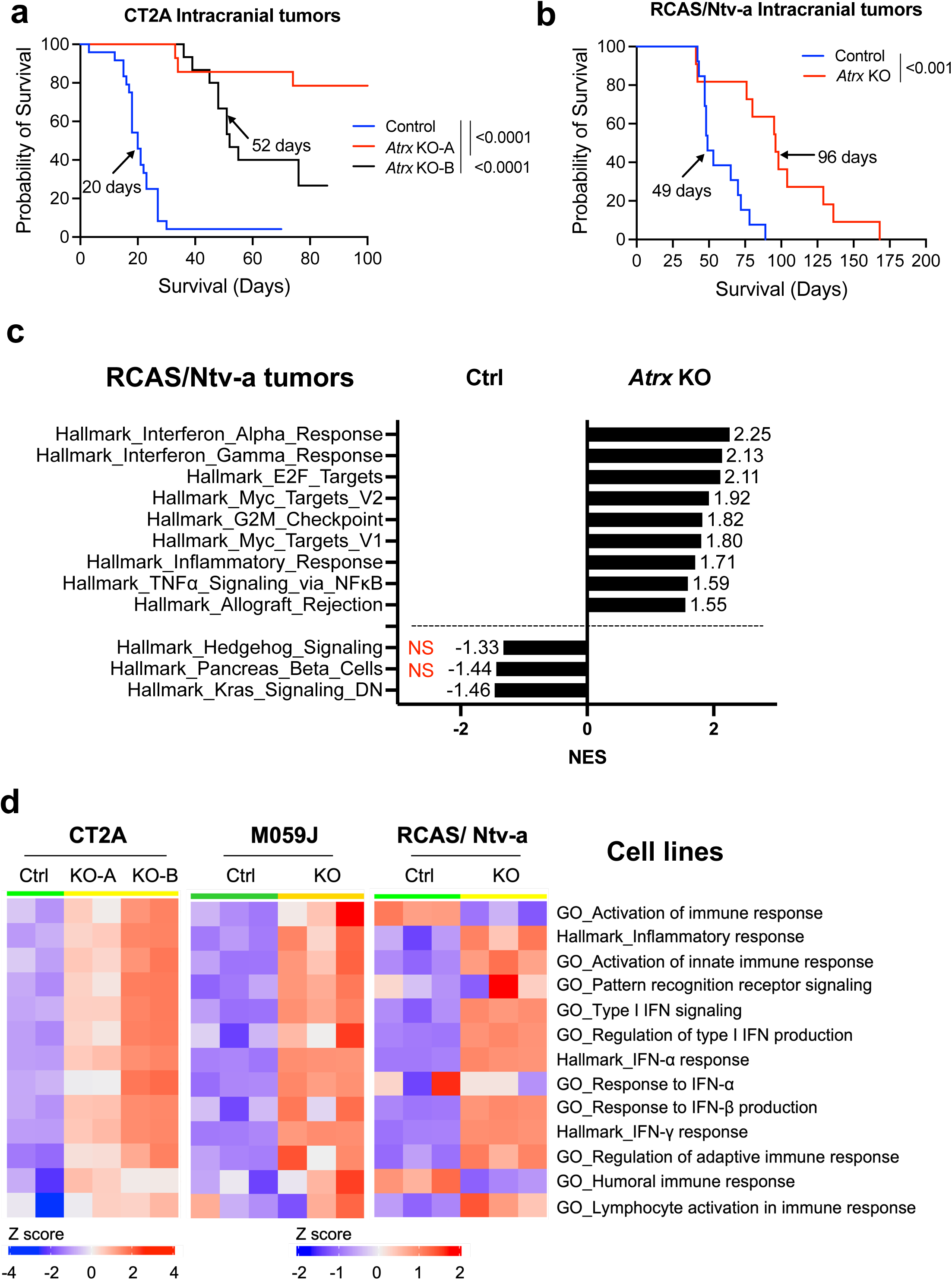
ATRX depletion is associated with improved survival *in vivo*. (**a**) Kaplan Meir survival curves for C57BL/6 mice bearing intracranial CT2A CRISPR ctrl (Control) (n=24), *Atrx*-KO clones, KO-A (n=14) and KO-B (n=15) tumors. Median survival is indicated in days for each group except Atrx KO-A group. P-values representing group comparisons were calculated using log-rank test. (**b**) Kaplan Meir survival curves for C57BL/6 mice bearing intracranial RCAS/ Ntv-a *Atrx^+/+^* (Ctrl) (n=13) and *Atrx^fl/fl^* (KO) (n=11) tumors. P-values representing group comparisons were calculated using log-rank test. (**c**) GSEA of Hallmark gene sets from RCAS/ Ntv-a *Atrx*-KO tumors (n=3) compared to *Atrx*^WT^ (Ctrl) tumors (n=3). Normalized enrichment scores (NES) for gene sets that have adjusted p value <0.05 (except Hallmark_Hedgehog signaling and Hallmark_Pancreas Beta cells, highlighted in red) are shown. (**d**) Heatmap from ssGSEA showing enrichment for various innate immune related gene sets in CT2A *Atrx* KO-A and KO-B clones, M059J *ATRX*-KO cells and RCAS/Ntv-a *Atrx*-KO cells compared to their respective *ATRX*^WT^ counterparts. n=2 technical replicates for CT2A expression data; n= 3 technical replicates for M059J expression data; n=3 biological replicates (3 consecutive cell passages) for RCAS/Ntv-a cell line expression data.

To probe the molecular and cellular mechanisms underlying impaired *in vivo* growth in the ATRX-deficient context, we performed RNA-seq on explanted tumors from RCAS/Ntv-a model and cultured cell lines from CT2A, M059J and RCAS model systems. GSEA of RCAS/Ntv-a tumors demonstrated differential engagement of cell cycle pathways in ATRX-deficient models relative to ATRX-intact counterparts, implicating tumor cell autonomous molecular mechanisms (Fig. 2c). However, we also found increased GSEA correlations with a variety of immune signaling networks in ATRX-deficient models, including the GO terms, inflammatory signaling, PRR signaling, activation of innate immune response, type I interferon signaling and response to IFNα and β (Fig. 2d; Supplementary Fig. 4a-b). Analysis of differential expression and baseline protein levels of genes involved in innate immune signaling indicated increased expression of RIG-I, MDA-5 (gene name – *Ifih1*), STAT1 and ISG15 in both CT2A *Atrx*-KO clones, KO-A and KO-B and M059J *ATRX*-KO cells compared to *ATRX*^WT^ counterparts, while increased expression of *Ifih1* and *Stat1* was observed in RCAS/Ntv-a *Atrx*-KO cells (Fig. 3a–b; Supplementary Fig. 5a-b). Moreover, several other genes involved in immune regulation, including *Jak1, Irf7, Irf9* and chemokines like *Ccl2, Ccl5* and *Cxcl10* were induced in CT2A *Atrx*-KO clones, KO-A and KO-B, M059J *ATRX*-KO and RCAS/Ntv-a *Atrx*-KO cells (Fig. 3a; Supplementary Fig. 5a-b).

**Figure 3:**
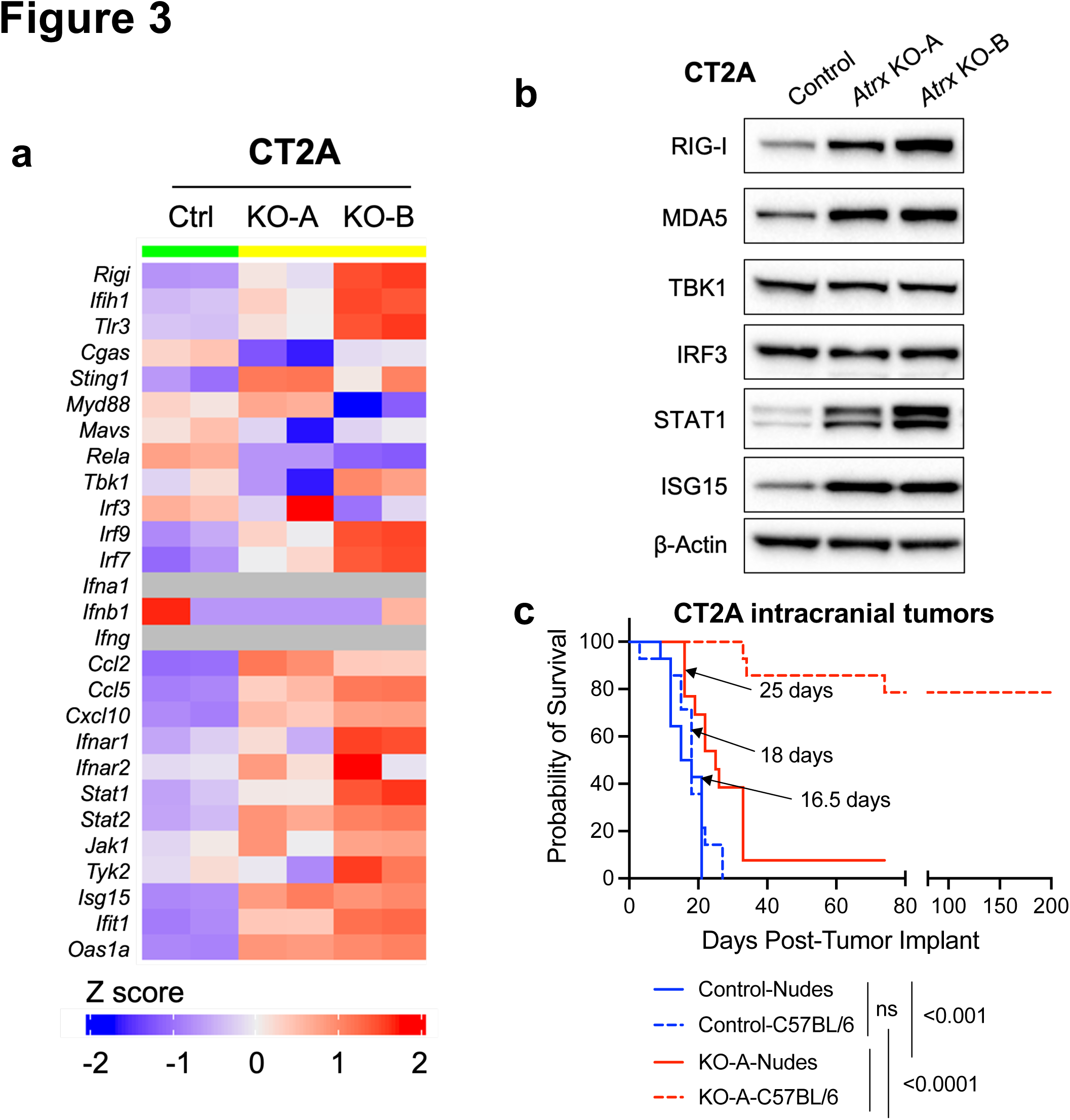
ATRX loss is associated with increased baseline inflammation *in vitro*. (**a**) Heatmap showing differential expression of immune-related genes in CT2A CRISPR control (Ctrl), *Atrx* KO-A and *Atrx* KO-B cells. n=2 technical replicates. (**b**) Western blot using lysates from CT2A CRISPR ctrl (Ctrl) and *Atrx*-KO clones, KO-A and KO-B screened for proteins involved in innate immune signaling. β-actin serves as the loading control. (**c**) Kaplan Meir survival curves for BALB/c nude and C57BL/6 mice bearing intracranial CT2A CRISPR ctrl (Control) or *Atrx* KO-A tumors. N values: Control-Nudes =14, Control-C57BL/6 =14, KO-A-Nudes =13, KO-A-C57BL/6 = 14. P-values represent group comparisons calculated using log-rank test.

To delineate the extent to which tumor cell-autonomous and immune microenvironmental factors impaired the growth of our ATRX-deficient glioma models *in vivo*, we compared survival among mice carrying *Atrx*^WT^ or *Atrx* KO-A tumors in either BALB/c nude mice, which lack cellular immunity, or C57BL/6 immune-intact contexts for our CT2A model. We found that differences in allograft growth between ATRX-intact and –deficient glioma were much less apparent in BALB/c nude hosts compared to C57BL/6 immunocompetent hosts (Fig. 3c). Taken together, these findings strongly suggest that the indolent growth of ATRX-deficient glioma models is largely attributable to immune microenvironmental effects.

### ATRX-deficient glioma models upregulate cytokine secretion and demonstrate increased immune cell infiltration

Having implicated the immune microenvironment as a potential causative factor promoting indolence in ATRX-deficient glioma, we sought to determine the extent to which ATRX depletion promotes increased immune cell infiltration in our *in vivo* models. To this end, we analyzed dissociated *Atrx*^WT^ and *Atrx*-KO CT2A tumor-bearing brain hemispheres from immunocompetent mice by flow cytometry (Fig. 4a). ATRX loss led to an overall significant increase in CD3+ T cells and, in particular, CD4+ T-cells within the tumor bearing hemispheres of allografted mice, while macrophage infiltration was reduced in the Atrx KO tumors. Moreover, RNA-seq analysis across CT2A, M059J, and RCAS/Ntv-a cell lines revealed upregulated transcripts for multiple chemokines/cytokines, including CCL2, CCL5, CXCL10 and IFNβ, in the ATRX-deficient context, potentially providing an underlying mechanism for immune cell recruitment (Fig. 3a; Supplementary Fig. 5a). We then confirmed upregulated cytokine secretion in ATRX-deficient isogenics for CT2A and RCAS/Ntv-a lines (Fig. 4b, c). Taken together, these findings mirror that of human tumors (Fig 1), where ATRX deficiency was associated immune cell infiltration and pro-inflammatory signaling in gliomas.

**Figure 4:**
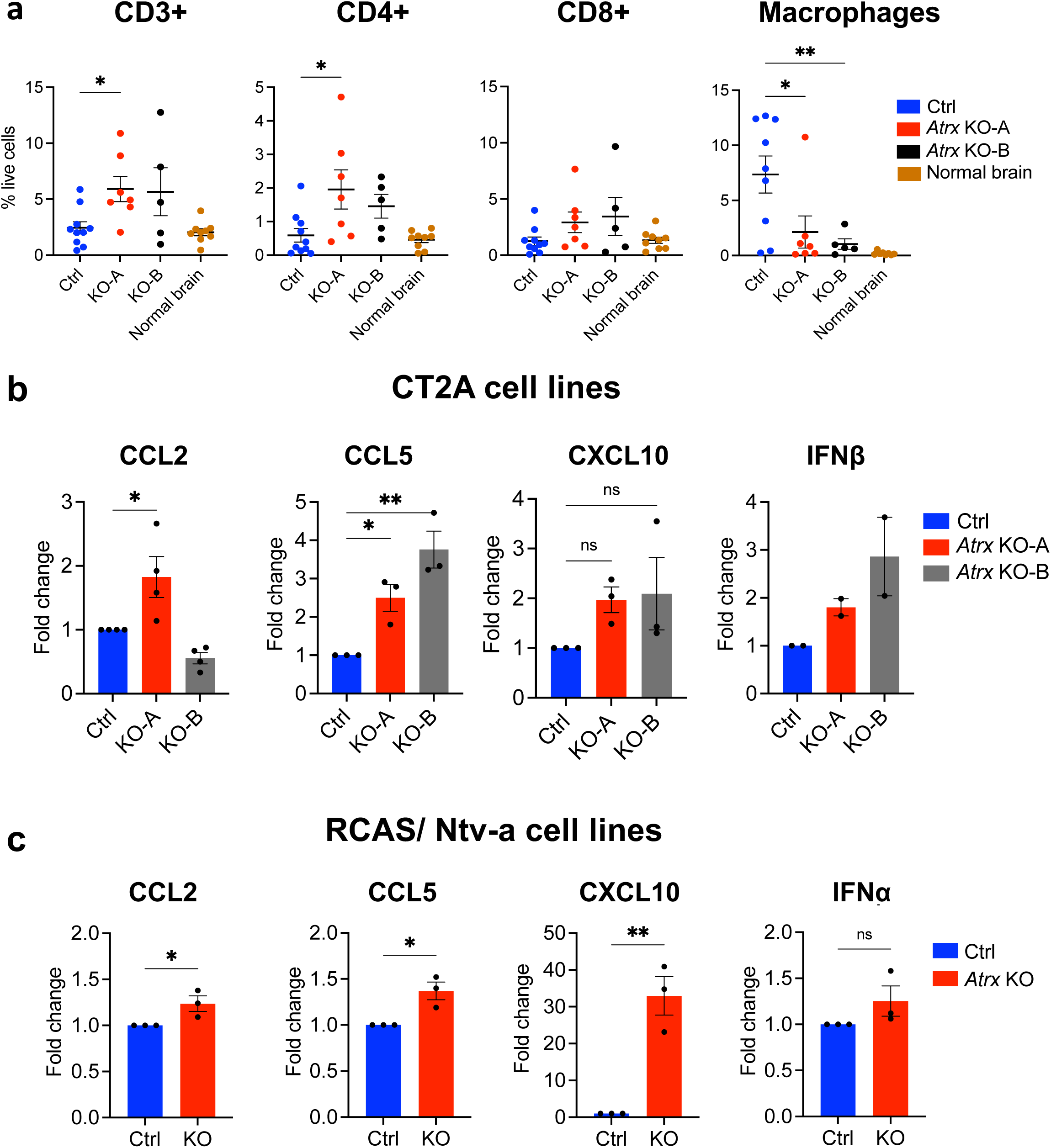
ATRX deficiency leads to increased T-cell infiltration & cytokine secretion. (**a**) Flow cytometry analysis of CT2A CRISPR control, *Atrx* KO-A or KO-B tumor-bearing hemispheres showing percent live cell density of CD3+, CD4+ and CD8+ T-cells and macrophages. n=10 for CRISPR control – CD3+, CD4+, CD8+ markers; n=9 for CRISPR control – macrophage markers; n=7 for *Atrx* KO-A – all markers; n=5 for *Atrx* KO-B – all markers; n=9 for normal brain – CD3+, CD4+, CD8+ markers; n=8 for normal brain – macrophage markers. Asterisks denote significant one-way ANOVA with Dunnett’s post-hoc test comparing CRISPR control with *Atrx* KO tumor-bearing hemispheres (*: p<0.05, **: p<0.01). (**b, c**) Cytokine levels in conditioned media from untreated CT2A CRISPR control (Ctrl), *Atrx* KO-A and KO-B clones (**b**), and RCAS/Ntv-a *Atrx^+/+^* (Ctrl) and *Atrx^-/-^* (KO) cell lines (**c**). Supernatant cytokines were analyzed by cytokine bead arrays for antiviral and proinflammatory cytokines. Data indicates fold change values normalized to CRISPR control (Ctrl; CT2A model) or *Atrx^+/+^* (Ctrl; RCAS/Ntv-a), shown as mean ± SEM. N values: CT2A model – biological replicates: n=4 for CCL2; n=3 for CCL5, CXCL10; n=2 for IFNβ; RCAS/Ntv-a model – technical replicates: n=3 for all cytokines. Asterisks denote significant one-way ANOVA with Dunnett’s post-hoc test (CT2A model) and significant unpaired, two-tailed t-test (RCAS/Ntv-a model) (*: p<0.05; **: p<0.01).

### ATRX depletion leads to enhanced innate responses to dsRNA agonists

As described above, *ATRX*-KO cell lines demonstrated increased expression of RIG-I and MDA5 dsRNA sensors. However, the extent to which ATRX loss influence dsRNA-triggered innate immune responses has not been explored. Treatment with either low molecular weight (LMW) or high molecular weight (HMW) poly(I:C)—a synthetic dsRNA analog— resulted in an augmented secretion of cytokines including CCL2, CCL5 and CXCL10 in CT2A *Atrx*-KO clones (KO-A – Fig. 5a; KO-B – Supplementary Fig. 7a), IFNβ, IL28α/β, IL29 and IL6 in M059J *ATRX*-KO cells (Fig. 5b); CXCL1, CCL2 and CXCL10 in RCAS/Ntv-a *Atrx*-KO cells (Fig. 5c). Poly(I:C) treatment also induced higher STAT1 phosphorylation and ISG15 expression, both markers of innate immune signaling, in all three ATRX-deficient isogenic models. Implying this effect is due to enhanced sensing of dsRNA, poly(I:C) strongly induced phosphorylation of the innate immune transcriptional regulator phosphorylated early during antiviral signaling, IRF3, in *Atrx*/*ATRX*-KO CT2A and M059J cells (Fig. 5d–f; Supplementary Fig. 7b). These observations suggest that ATRX inactivation sensitizes glioma cells to dsRNA.

**Figure 5:**
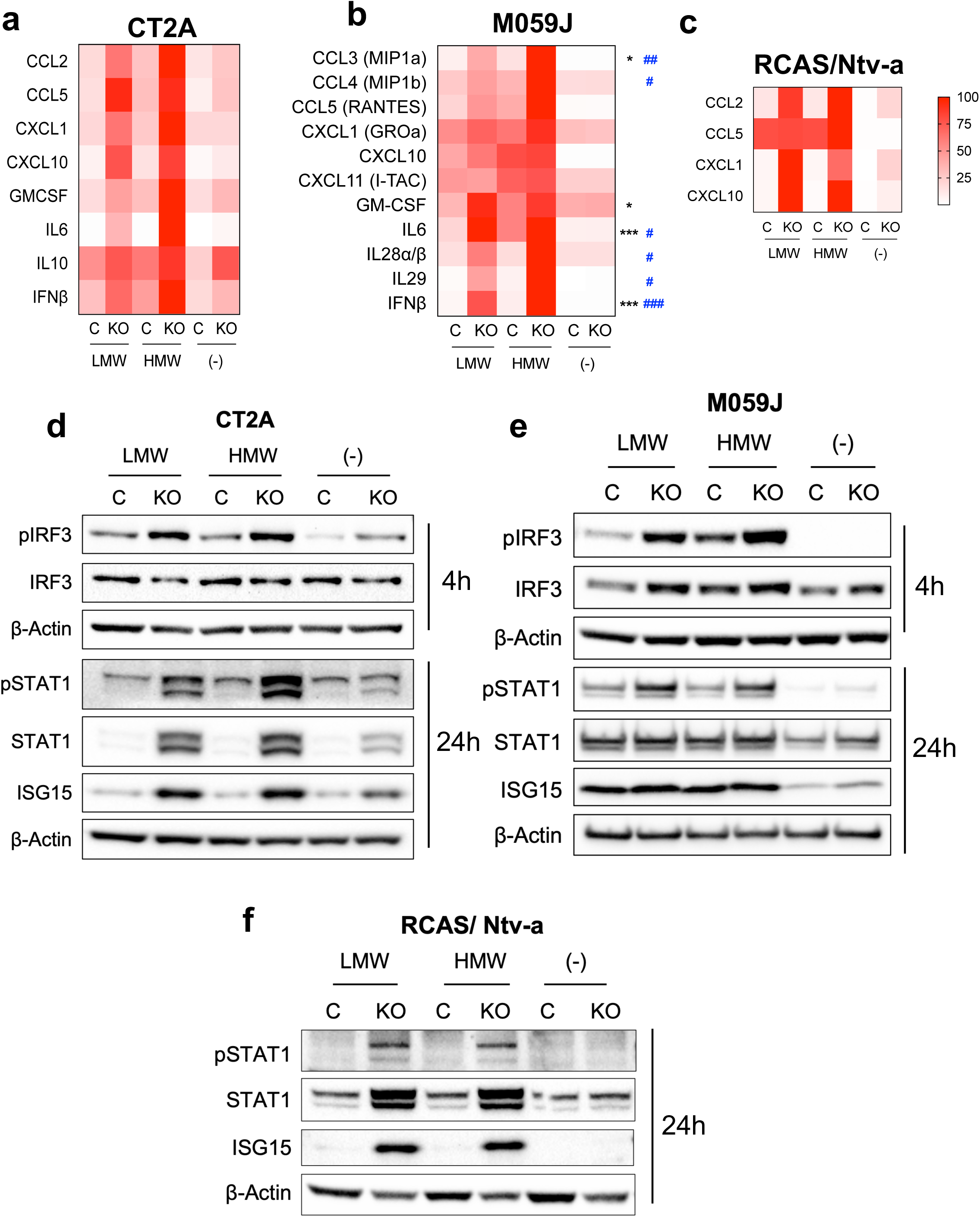
ATRX depletion sensitizes cells to poly(I:C), a dsRNA agonist. (**a, b, c**) Cytokine levels in conditioned media from CT2A CRISPR control and *Atrx* KO-A cells (**a**), M059J CRISPR control and *ATRX*-KO cells (**b**) and RCAS/Ntv-a *Atrx^+/+^* or *Atrx^-/-^* cell lines (**c**) treated with 10μg/ml poly(I:C) LMW or HMW for 24hrs. Supernatant cytokines were analyzed by cytokine bead arrays for antiviral and proinflammatory cytokines. n=2 biological replicates for CT2A; n=2 biological replicates for M059J; n=1 for RCAS/Ntv-a cell lines. For heatmap generation, maximum values for each cytokine were set to 100%. Asterisks & hashtags denote significant one-way ANOVA with Sidak’s post-hoc test comparing both poly(I:C) LMW (*) and poly(I:C) HMW (#) between *ATRX*-KO and CRISPR control cell lines (*, #: p<0.05; **, ##: p<0.01; ***, ###: p<0.0001). Only cytokines and signaling proteins with observed induction after treatment with poly(I:C) are included. (**d, e, f**) Western blot using lysates from CT2A CRISPR control and *Atrx* KO-A cells (**d**), M059J CRISPR control and *ATRX*-KO cells (**e**) and RCAS/Ntv-a *Atrx^+/+^* or *Atrx ^-/-^* cells (**f**) treated with poly(I:C) LMW or HMW for 2hrs (M059J), 4hrs (CT2A) or 24hrs CT2A, M059J, RCAS/Ntv-a), screened for pIRF3/IRF3, pSTAT1/STAT1 and ISG15 involved in innate immune signaling. β-actin serves as the loading control.

### Co-expression of IDH1^R132H^ with ATRX loss attenuates baseline innate immune signaling in gliomas

Recent work has shown that mutant *IDH1* cooperates with ATRX loss in promoting the alternative lengthening of telomere (ALT) phenotype in gliomas, suggesting an interplay between the two molecular abnormalities^31^. Furthermore, mutant *IDH1* has been associated with an immunosuppressive phenotype characterized by decreased tumor infiltration by T-cells and reduced expression of cytotoxic T lymphocyte-associated genes, pro-inflammatory cytokines, and chemokines^25, 32^. These phenotypes, along with many others, has been linked to production of the oncometabolite D-2-hydroxyglutarate (D2HG) by mutant *IDH*^33, 34^. To investigate the impact of co-occurring *IDH1* and *ATRX* mutations on innate immune signaling in glioma, we generated CT2A cells expressing mutant IDH1^R132H^ in both *Atrx*^WT^ and *Atrx*-KO genetic backgrounds. IDH1^R132H^ expression and D2HG production were both confirmed in relevant cell lines by immunoblotting and mass spectrometry, respectively (Fig. 6a, b). Expression of IDH1^R132H^ did not alter *in vitro* proliferation, regardless of *ATRX* status (Fig. 6c).

**Figure 6:**
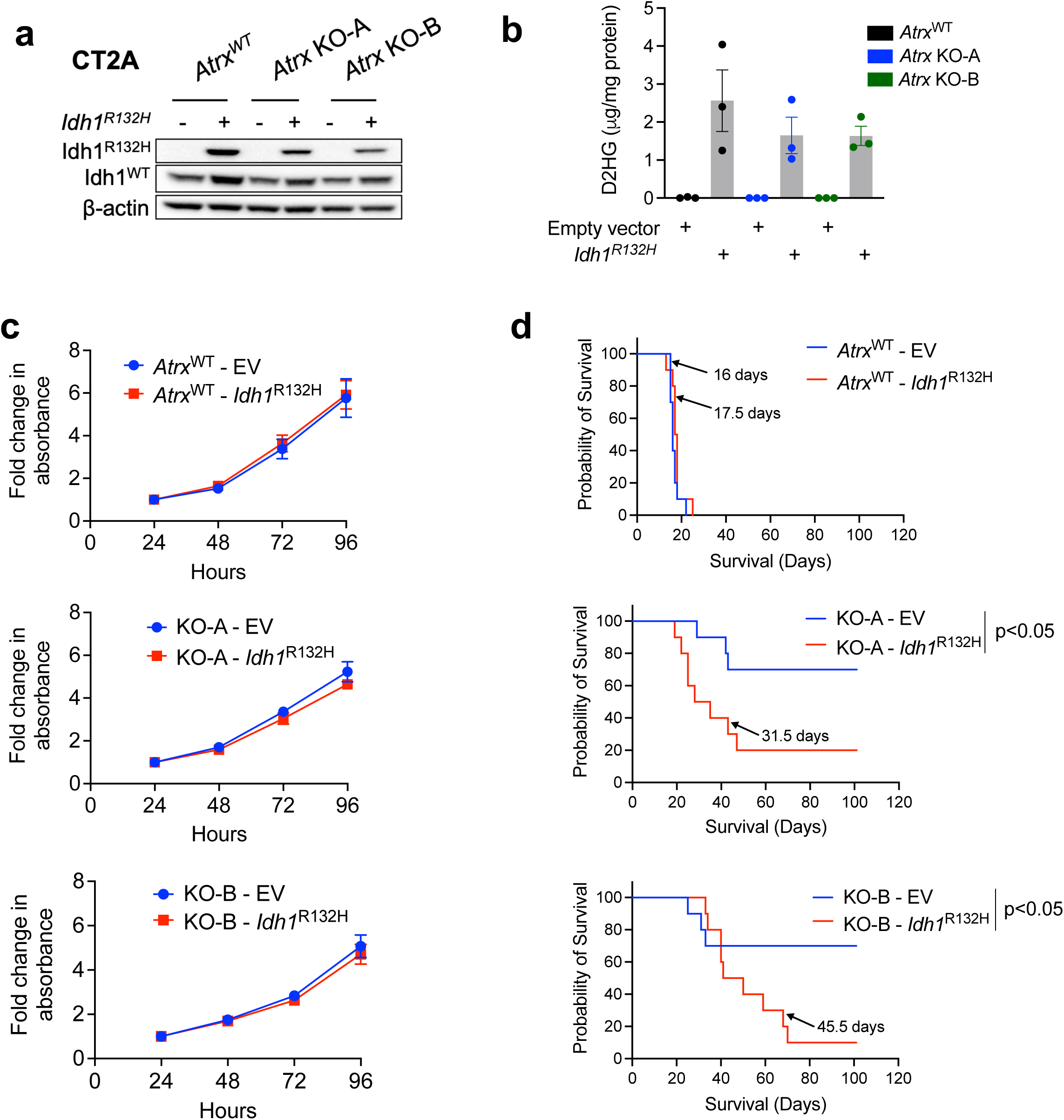
Generation & characterization of IDH1^R132H^-expressing CT2A cells. (**a**) Lysates from CT2A CRISPR control (*Atrx*^WT^) or *Atrx*-KO cells (KO-A, KO-B) expressing exogenous IDH1^R132H,^ or empty vector were screened by immunoblotting for IDH1^R132H^ and wildtype IDH1 expression. β-actin serves as the loading control. (**b**) D2HG levels in cell pellets from CT2A CRISPR control (*Atrx*^WT^) or *Atrx*-KO cells (KO-A, KO-B) expressing IDH1^R132H^ or empty vector in culture for 72hrs, normalized to total protein. n=3 biological replicates. (**c**) Growth assays of CT2A CRISPR control (*Atrx^WT^*) or *Atrx*-KO cells (KO-A, KO-B) expressing IDH1^R132H^ or empty vector in culture for 96 hrs. Data represents mean ± SEM. n= 3 biological replicates. (**d**) Kaplan Meir survival curves for C57BL/6 mice bearing intracranial CT2A CRISPR ctrl-empty vector control (*Atrx*^WT^ – EV), CRISPR ctrl-*Idh1*^R132H^ (*Atrx*^WT^ – *Idh1*^R132H^), *Atrx* KO-A – empty vector control (KO-A – EV), *Atrx* KO-A – *Idh1*^R132H^ (KO-A – *Idh1*^R132H^), *Atrx* KO-B – empty vector control (KO-B – EV) and *Atrx* KO-B – *Idh1*^R132H^ (KO-B – *Idh1*^R132H^) tumors. n=10 for each group. P-values represent group comparisons calculated using the log-rank test.

To determine the effect of IDH1^R132H^ co-expression on in vivo tumor growth, *Atrx*^WT^ and *Atrx*-KO CT2A cell lines, with or without *Idh1^R132H^* were intracranially allografted into immunocompetent mice. Resulting tumors exhibited largely identical histopathology, regardless of their underlying genetics (Supplementary fig. 8). However, we found that co-expression of IDH1^R132H^ shortened overall survival only in the *Atrx* KO context (Fig. 6d), suggesting that *IDH1^R132H^*contributes to glioma progression in *Atrx* deficient gliomas.

RNAseq analysis of our isogenic CT2A lines indicated that while ATRX deficiency upregulated various innate and adaptive immune-related gene sets, co-expression of IDH1^R132H^ in ATRX-deficient cells mitigated these effects (Fig. 7a; Supplementary Fig. 9, 10). For instance, differentially expressed genes involved in innate immune signaling like *Ddx58, Ifih1, Tlr3, Myd, Irf7, Stat1, Isg15*, enriched in *Atrx*-KO cells, were downregulated in *Atrx-*KO/ *Idh1^R132H^* cells (Fig. 7b; Supplementary Fig. 11a). IDH1^R132H^ expression was also associated with decreased baseline levels of key innate immune pathway proteins like RIG-I, MDA5, STAT1 and ISG15 in both the *Atrx*^WT^ and *Atrx*-KO contexts, although expression of IRF3 was unaffected across cell lines (KO-A – Fig. 7c; KO-B – Supplementary Fig. 11b). Moreover, secretion of cytokines like CCL2, CCL5, CXCL10 and IFNβ was similarly downregulated upon IDH1^R132H^ expression (KO-A – Fig. 7d; KO-B – Supplementary Fig. 11c). These findings indicate that co-existent IDH1^R132H^ tempers ATRX-deficient inflammatory signaling in gliomas, further supporting the notion that mutant *IDH1* confers an immunosuppressive phenotype and pointing to a pathogenically relevant immunological interplay between the defining molecular alterations of *IDH-*mutant astrocytoma.

**Figure 7:**
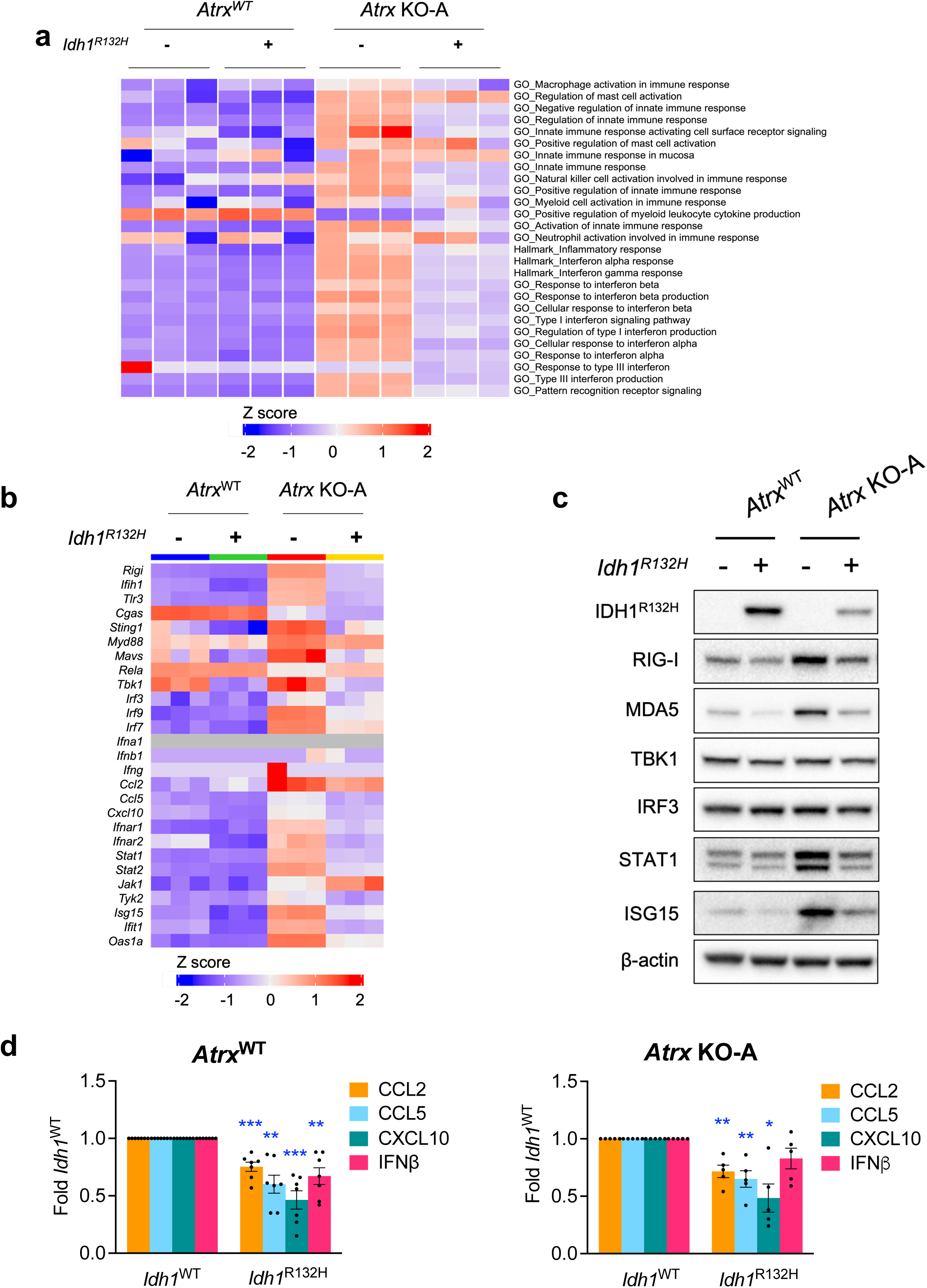
Idh1^R132H^ co-expression in Atrx-deficient cells dampens baseline innate gene expression. (**a**) Heatmap from ssGSEA showing loss of enrichment of various innate immune-related GO terms in *Atrx* KO-A/ *Idh1^R132H^* cells compared to *Atrx KO*-A/ *Idh1^WT^* cells. RNA was isolated from cells cultured for 72hrs. n=3 technical replicates per cell line. (**b**) Heatmap showing differential expression of immune-related genes in CT2A *Atrx*^WT^ and *Atrx* KO-A cells with or without *Idh1^R132H^*. n=3 technical replicates per cell line. (**c**) Lysates from CT2A CRISPR control (*Atrx*^WT^) or *Atrx* KO-A cells expressing exogenous IDH1^R132H,^ or empty vector cultured for 72hrs, were screened by Western blotting for various innate immune proteins. b-actin serves as the loading control. (**d**) Conditioned media from CT2A CRISPR control (*Atrx*^WT^) or *Atrx* KO cells (KO-A) cells expressing exogenous IDH1^R132H^, or empty vector cultured for 72hrs was assayed for antiviral and proinflammatory cytokines using a Legendplex assay kit. Data indicates fold change values normalized to corresponding *Idh1*^WT^ sample, shown as mean ± SEM. n= 7 biological replicates for *Atrx*^WT^ (CRISPR control) lines; n=5 biological replicates for *Atrx* KO-A lines. Asterisks denote significant two-tailed t-tests. (*: p<0.05; **: p<0.01; ***: p<0.001).

**Figure 8:**
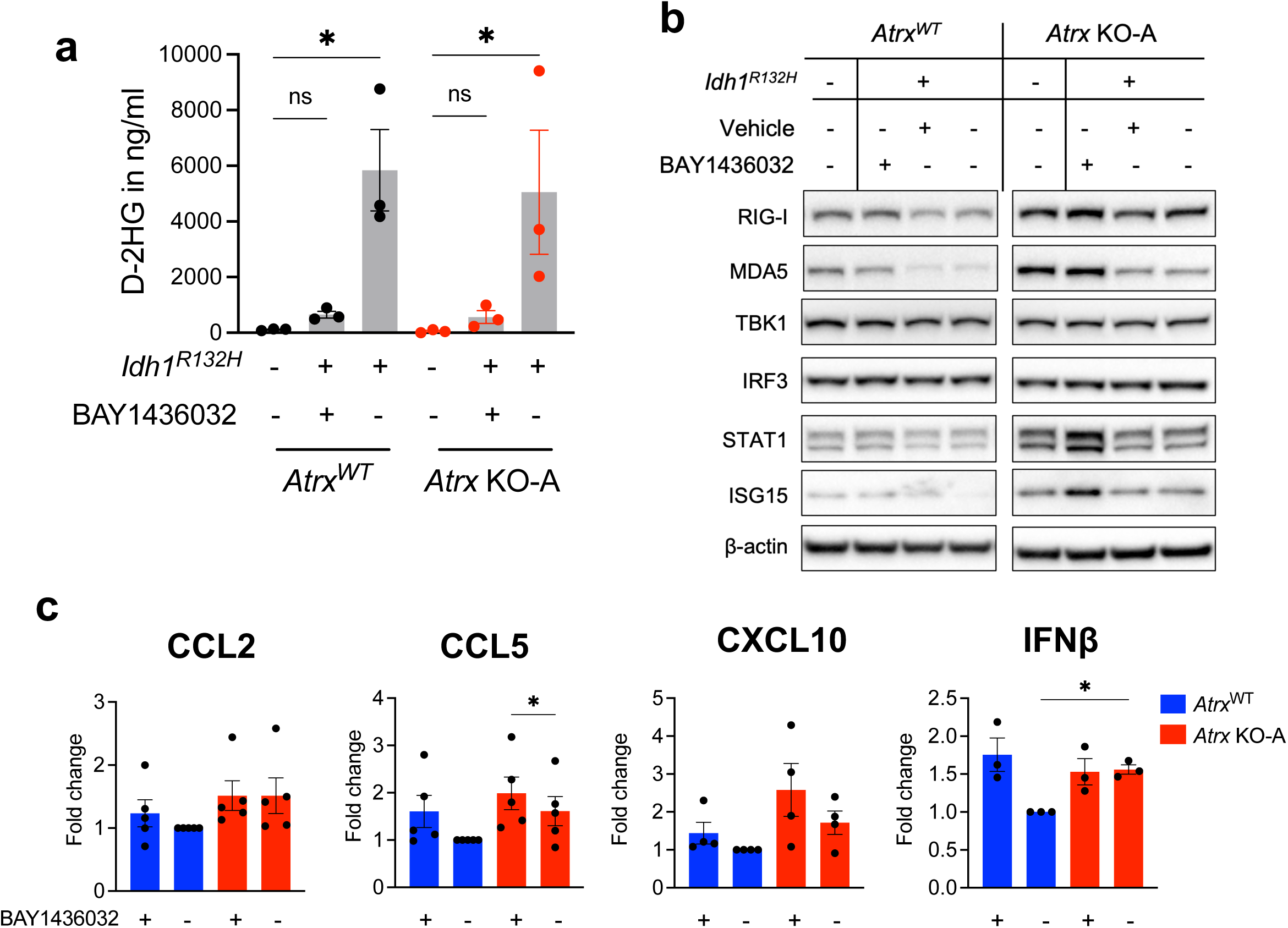
BAY1436032 partially reverses *IDH1^R132H^*-mediated immunosuppression. (**a**) D2HG levels in conditioned media from CT2A CRISPR control (*Atrx*^WT^) or *Atrx* KO-A cells expressing IDH1^R132H^, or empty vector treated with 1mM BAY1436032 or vehicle every day for 3 days, normalized to total protein. n=3 biological replicates. Asterisks indicate significant p-values from one-way ANOVA with Sidak post-hoc test. (*: p<0.05). (**b**) Western blot using lysates from CT2A CRISPR control (*Atrx*^WT^) or *Atrx* KO-A cells expressing IDH1^R132H^ or empty vector that were treated with 1μM BAY1436032 or vehicle every day for 3 days, screened for proteins involved in innate immune signaling. β-actin serves as the loading control. (**c**) Cytokine/ chemokine levels in conditioned media from CT2A CRISPR control (*Atrx*^WT^) and *Atrx* KO-A cells expressing IDH1^R132H^ treated with 1μM BAY1436032 or vehicle every day for 3 days. Supernatant cytokines were analyzed by cytokine bead arrays for antiviral and proinflammatory cytokines. Fold change values normalized to vehicle treated *Atrx*^WT^ sample are shown as mean ± SEM. Biological replicates: n=5 for CCL2, CCL5; n=4 for CXCL10; n=3 for IFNβ. One-way ANOVA with Tukey’s post-hoc test did not reveal any significant differences between groups.

### Mutant *IDH1* inhibition partially reverses the immunosuppressive effects of *IDH1^R132H^*

Several mutant *IDH* inhibitors are currently being tested for safety and efficacy in the treatment of *IDH1*-mutant gliomas and acute myeloid leukemias, including BAY1436032, a pan-IDH1 inhibitor^35–37^. To evaluate the impact of IDH1^R132H^ inhibition on the immunological phenotypes detailed above, we treated our panel of CT2A isogenic cell lines with 1μM BAY1436032, leading to reduced D2HG levels in both the *Atrx*^WT^ and *Atrx*-KO cell lines (KO-A – Fig. 8a; KO-B – Supplementary Fig. 12a). Treating IDH1^R132H^-expresing cells with BAY1436032 for 72hrs increased baseline expression of RIG-I, MDA5, STAT1 and ISG15, irrespective of the *Atrx* status (KO-A – Fig. 8b; KO-B – Supplementary fig. 12b), reverting levels to those seen in the *Idh1*^WT^ context and demonstrating that crucial components of mutant *IDH1*-associated immunomodulation are reversible. BAY1436032 treatment did not affect expression of these innate immune proteins in *Idh1*^WT^ cells (data not shown). Interestingly, BAY1436032 also induced pro-inflammatory cytokines, including CCL2, CCL5, CXCL10 and IFNβ in *Atrx*^WT^*/ Idh1*^R132H^ cells to levels equivalent to those of untreated *Atrx-*KO*/ Idh1*^R132H^ cells (KO-A – Fig. 8c; KO-B – Supplementary Fig. 12c). However, by contrast, BAY1436032 treatment of *Atrx-*KO*/ Idh1*^R132H^ cells failed to significantly increase cytokine secretion, though CCL5 and CXCL10 levels were non-significantly elevated. Thus, the immunomodulatory phenotype induced by mutant *IDH1* in the context of ATRX deficiency is partially reversible with inhibitory therapy.

### *Atrx*-KO/ *Idh1*^R132H^ cells retain sensitivity to dsRNA-based immune agonists

To determine the extent to which ATRX-deficient glioma models remain sensitive to dsRNA immune agonism in the context of *IDH1^R132H^*, we subjected our isogenic CT2A lines to HMW poly(I:C), monitoring levels of key pathway constituents. Poly(I:C) treatment effectively induced IRF3 and STAT1 phosphorylation and ISG15 expression in IDH1^R132H^-expressing cells with concurrent ATRX deficiency (KO-A – Fig. 9a; KO-B – Supplementary Fig. 13a). While pIRF3 induction was equivalent in both *Atrx* KO-A and *Atrx* KO-B lines expressing IDH1^WT^ or IDH1^R132H^, pSTAT1 and ISG15 induction was somewhat weakened in *Atrx* KO-A*/ Idh1*^R132H^ cells. Nevertheless, pSTAT1 and ISG15 were induced at equivalent levels in both *Atrx* KO-B*/ Idh1*^WT^ and *Atrx* KO-B*/ Idh1*^R132H^ lines. Poly(I:C) treatment also increased secretion of cytokines like CXCL1, CCL2, CCL5, CXCL10 and IL6 in *Atrx* KO lines compared to *Atrx*^WT^ lines, irrespective of the *Idh1* status (data not shown). Moreover, the presence of IDH1^R132H^ did not impact cytokine secretion in either *Atrx* KO-A or *Atrx* KO-B lines (Fig. 9b; Supplementary Fig. 13b), consistent with the trends observed for pIRF3, pSTAT1, and ISG15 described above. These results indicate that IDH1^R132H^ co-expression in ATRX-depleted cells does not substantively impair dsRNA-mediated induction of innate immune response. Accordingly, these data further support the notion that the agonizable state of innate immune signaling in ATRX-deficient glioma exists largely independent of *IDH1* mutational status.

**Figure 9:**
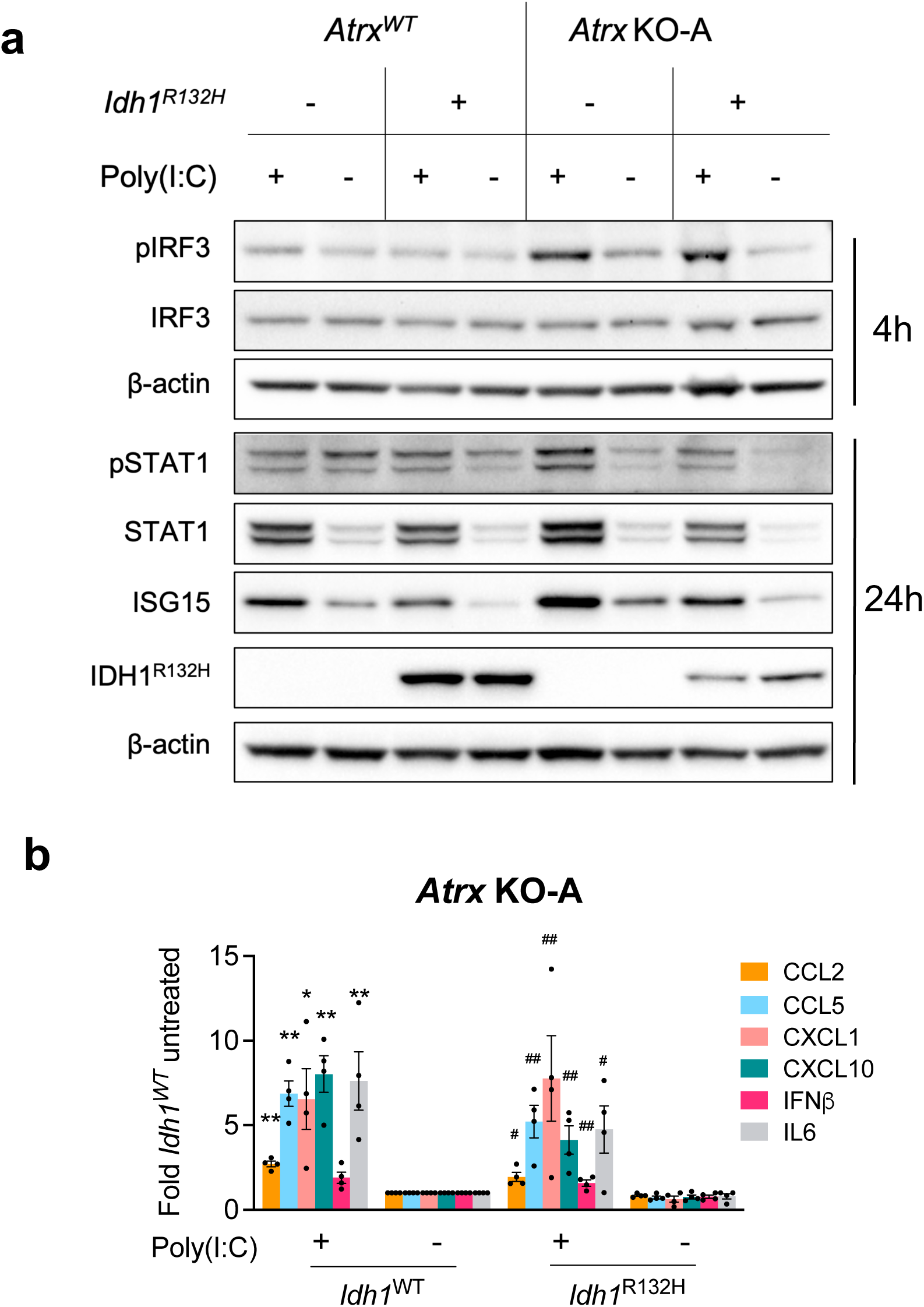
*Atrx* KO/ *Idh1^R132H^* cells retain sensitivity to poly(I:C). (**a**) Western blot using lysates from CT2A CRISPR control (*Atrx*^WT^) or *Atrx* KO-A cells expressing IDH1^R132H^ or empty vector that were treated with 10mg/ml poly(I:C) HMW for 4hrs or 24hrs and screened for proteins involved in innate immune signaling. **(b)** Cytokine levels in conditioned media from CT2A *Atrx* KO-A cells expressing IDH1^R132H^ or empty vector treated with poly(I:C) HMW for 24hrs. Supernatant cytokines were analyzed by cytokine bead arrays for antiviral and proinflammatory cytokines. Fold change values normalized to untreated *Atrx* KO-A/ *Idh1*^WT^ sample are shown as mean ± SEM. n= 4 biological replicates. Asterisks and hashtags indicate significant p-values from one-way ANOVA with Tukey’s post hoc test comparing poly(I:C) HMW with untreated samples for *Atrx*-KO-A – EV control line (*) or *Atrx*-KO-A – *Idh1*^R132H^ (*, #: p<0.05; **, ##: p<0.01; ***, ###: p<0.0001).

**Figure 10:**
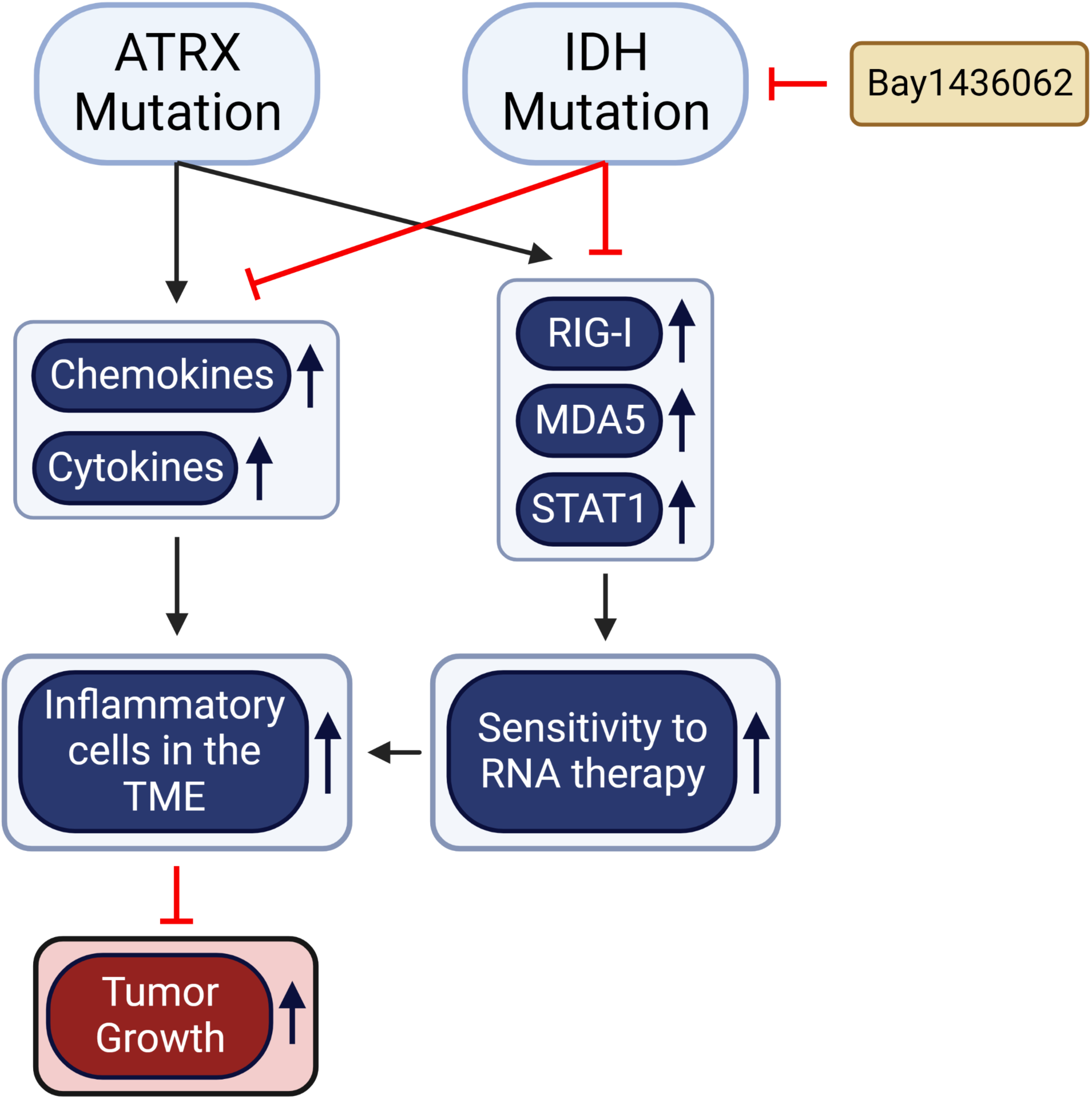
Cartoon summarizing the effect of *ATRX* mutations on baseline and dsRNA-mediated innate signaling and its interplay with *IDH* mutations in gliomas.

## Discussion

Leveraging the immune system in the context of gliomas has been met with relatively diminished return in comparison to its considerable promise in other cancers. This is partially due to the immunosuppressive microenvironment, which has led to tremendous efforts aimed at identifying novel routes for augmenting immune activity and response^38^.

While the association between mutations in *ATRX* and *IDH1* has been known for over a decade, the details of their interaction and basis for recurrent co-occurrence in astrocytoma remains unclear. Furthermore, the implication of these mutations on immune-therapeutic responsiveness is understudied. Here we describe the complex and contrasting impact of these disease-defining molecular alterations on the immune microenvironment and characterize strategies for therapeutic targeting (Fig. 10).

Mutations in *IDH* contribute to the overall immunosuppressive microenvironment characteristic of gliomas^25, 32, 39^. However, the precise role of molecular alterations co-occurring with *IDH* mutations, including *ATRX* inactivation, are beginning to be elucidated ^23–26, 32, 39^. We demonstrated that *ATRX* mutant astrocytomas display a more pro-inflammatory gene expression profile compared to oligodendroglioma counterparts. Suggesting a connection between ATRX loss and immune-modulatory behavior, pro-inflammatory gene expression profiles are enriched among *ATRX* mutant low-grade gliomas. This was apparent through analysis of both TCGA-LGG data and RNAseq analysis of our genetically engineered murine and human glioma models. In the process of developing our orthotopic models (both CT2A and RCAS/Ntv-a), we observed prolonged survival among animals bearing ATRX*-*inactivated tumors, associated with an increase in T-cell infiltration, enhanced innate immune signaling, and production of cytokines and chemokines indicative of an activated immune response *in vitro*. Interestingly, this survival phenotype was observed to a lesser extent when ATRX-deficient cells were implanted into nude animals, implicating immune microenvironmental engagement as a fundamental driver of this phenotype. The genetic and expression data of ATRX-deficient human and murine systems, coupled with the *in vitro* upregulation of RIG-I and MDA5, and increased cytokine production of CCL2, CCL5, CXCL10, and IFNβ, depicted a robust pro-inflammatory phenotype and stimulation of the innate arm of the immune system. In total, these findings link ATRX loss with a pro-inflammatory state.

This observed induction, suggested an enhanced dsRNA sensing mechanism that exposes a therapeutic vulnerability of *ATRX* mutated cells to dsRNA agonists. To this end, we investigated the response of ATRX-deficient cells to the dsRNA agonist poly(I:C), which led to secretion of cytokines indicative of immune stimulation. While this phenotype was observed across murine and human models, the extent of *in vivo* response remains to be seen and will be the foundation of future studies. Specifically, understanding how dsRNA agonists can mount an enhanced immune response and whether this confers a therapeutic benefit under *in vivo* conditions will be paramount. Indeed, various treatment approaches, including oncolytic virotherapies grounded in the induction of RNA sensing and stimulation of the innate immune system, have shown promise for several cancers including GBM^11, 12, 14^. For example, recent advances utilizing the recombinant nonpathogenic polio-rhinovirus chimera (PVSRIPO) have shown an improved survival rate among treated patients with recurrent glioblastoma^11^.

Considering their frequent co-occurrence as well as their respective roles in immune response, understanding how *ATRX* and *IDH* mutations influence immune state in combination is warranted. We demonstrated that among ATRX-deficient cell models, the presence of the *IDH1* mutation has negligible impact on proliferation; however, upon intracranial implantation, mutant *IDH1* significantly increases tumor aggressiveness, as indicated through a shorter survival and greater tumor penetrance. Interestingly, while mutant *IDH1* did not impact the enrichment of various innate immune-related gene sets in the context of wildtype *ATRX*, it did depress the induction of the innate immune response when ATRX was lost. Similarly, many of these genes were downregulated in response to mutant *IDH1*, reverting their expression to the baseline levels seen in *ATRX*^WT^ cells, conferring an overall immunosuppressive impact, and dampening of the pro-inflammatory effect of ATRX loss. Nevertheless, despite this mutant *IDH1*-mediated immunosuppressive phenotype, ATRX-deficient cells remained capable of mounting a robust innate response upon poly(I:C) treatment, a key finding of therapeutic relevance.

The impact of *IDH* mutation and ATRX deficiency on immune function remains an open question in the literature, explored in multiple recent publications. One earlier study found that ATRX loss in the context of *IDH1* and *TP53* mutations confers an immunosuppressive state characterized by altered cytokine secretion, leading to increased ATRX-deficient tumor growth^24^. Of note, increased CCL2 and CCL5 levels were observed in both our study and this prior report, while expression of other cytokines differed, possibly owing to distinctions in model background. It is also worth noting that the *in vivo* arm of that study relied on GL261, a model known for high baseline immunogenicity^40^. In contrast, both our CRISPR and RCAS/Ntv-a models displayed low baseline immunogenicity, better reflecting the human context. Another group developed murine glioma models based on *Atrx* KO followed by overexpression of mutant *Idh1* and described an upregulation of pro-inflammatory gene regulatory programs as characterized by single cell RNA-seq and ATAC-seq^23^. However, accumulation of immunosuppressive M2 macrophages was also observed in this model, complicating their overall interpretations of ATRX loss and its impact. Of note, both preceding studies examining the immunologic impact of ATRX deficiency relied heavily on RNA-level data, while our work integrated protein and gene set level data, along with functional changes *in vivo*. Importantly, our study adds to the prior work on this topic by demonstrating the hypersensitivity of ATRX deficient cells to innate immune agonism, and by demonstrating the ability of mutant *IDH* inhibition to unmask the pro-inflammatory consequences of ATRX loss. These findings are further supported by our analysis of human TCGA data, showing that combined ATRX loss and IDH mutation in astrocytomas yields a pro-inflammatory phenotype, relative to *IDH* mutation alone.

In light of the mitigating effects conferred by *IDH* mutation on the pro-inflammatory phenotype of *ATRX* deficiency, we considered the possibility that mutant *IDH1* inhibition could restore immune pathway engagement in our experimental model. Indeed, when the pan-mutant *IDH1* inhibitor, BAY1436032^35–37^ was administered to *Atrx-KO*/*Idh1*^R132H^ lines, we found increased expression of RIG-I, MDA5, STAT1, and ISG15, relative to vehicle-treated controls. These trends suggest that mutant *IDH1*-mediated suppression of innate immune response in ATRX deficient lines can be at least partially rescued by BAY1436032, and this strategy might also potentiate therapeutics seeking to exploit the ATRX-deficient pro-inflammatory phenotype.

## Methods

### Cell lines

Human M059J cells were obtained from ATCC (Manassas, VA). Mouse CT2A cells were a kind gift of Peter Fecci (Duke University Medical Center). Human 293FT cells were obtained from Thermo Scientific (Waltham, MA). M059J cells were maintained in DMEM/F12 medium containing 2.5mM L-glutamine, 15mM HEPES, 0.5mM sodium pyruvate, 1.2g/L sodium bicarbonate and 10% fetal bovine serum (FBS). CT2A and 293FT cells were maintained in DMEM (high glucose) containing 10% FBS. All cell lines were cultured without antibiotics. Cell lines were routinely tested for mycoplasma contamination using the MycoAlert^®^ kit (Promega, Madison, WI).

### Generation of CRISPR-Cas9-mediated *ATRX* KO cell lines

For generating *ATRX* KO cell lines, single guide RNAs (sgRNAs) targeting exon 6 and exon 9 of human *ATRX* or mouse *Atrx* (Supplementary Table 1) were cloned into CRISPR-Cas9 lentiCRISPR v2 vector (gift from Feng Zhang, Addgene plasmid #52961). Lentivirus was produced in 293FT cells. Each cell line was transduced with a pair of lentiviruses targeting exon 6 and exon 9 simultaneously to ensure efficient gene knockout. Two days after transduction, cells were selected with puromycin for 5-7 days, and gene knockout confirmed by sequencing and western blot. Alternatively, lines were subjected to single cell cloning. A polyclonal population of *ATRX*-KO M059J cells and single cell clones isolated from CT2A *Atrx*-KO cells were used for experiments. ATRX knockout in these final reagents was confirmed by sequencing and immunoblotting. Since ATRX loss is associated with DNA damage and genomic instability^41^, only early passage *ATRX*-KO cell lines were used for experiments.

### Generation of *Idh1^R132H^* cell lines

IDH1^R132H^-expressing CT2A lines were generated using a retroviral MSCV-*Idh1*^R132H^ construct, or its corresponding empty vector control. Lentiviruses produced in 293FT cells were transduced into CT2A CRISPR control and *Atrx* KO clones, KO-A and KO-B lines. All cells were transduced twice to improve efficiency. Cells were allowed to expand for a week after transduction and expression of IDH1^R132H^ was confirmed by immunoblotting and immunofluorescence.

### *In vivo* CT2A model

Female BALB/c (JAX# 002019) or C57BL/6 mice (JAX #000664) aged 8 to 10 weeks were purchased from Jackson Laboratories (Bar Harbor, ME). All animal studies were performed in accordance with Duke Institutional Animal Care and Use Committee (IACUC) guidelines. CT2A cells were intracranially implanted into the right caudate nucleus (Implant coordinates: – 2.0, 0.5, 3.7/ 3.4) using an inoculation volume of 5μl. Mice were implanted with an inoculum of 150,000 cells suspended in 3% methylcellulose (30,000 cells/μl). Animals were monitored daily and euthanized upon development of neurological symptoms or signs of distress. Brains from all animals were harvested at necropsy. Brains were fixed overnight in 10% formalin, paraffin-embedded and sectioned (5μM thickness) for hematoxylin-eosin (H&E) and IDH1^R132H^ immunostaining.

### RCAS model – *in vivo* and *in vitro*

All animal experiments were performed in accordance with protocols approved by the MD Anderson Cancer Center (MDACC) Institutional Animal Care and Use Committees. *Nestin-tva mice* were generously provided by Dr. Eric Holland and ATRX-floxed mice were acquired from Dr. David Picketts. These mice were crossed, and pups genotyped by PCR from tail-derived genomic DNA. The RCAS-PDGFα-shp53 and RCAS-cre constructs was generously provided by Dr. Eric Holland. DF1 cells were transfected with the different RCAS retroviral plasmids using FuGENE 6 Transfection reagent (Promega, Cat. E2691), accordingly to the manufacturer’s protocol. For RCAS-mediated gliomagenesis, newborn mice were injected intracranially with 4×10^5^ DF1 cells per mouse, combining RCAS-PDGFa-shp53 and RCAS-Cre expressing cells at a 1:1 dilution. For xenografting experiments, adult mice were anaesthetized by 4% isofluorane and then injected with a stereotactic apparatus (Stoelting) as previously described^28^. After intracranial injection, mice were checked until they developed symptoms of disease (lethargy, poor grooming, weight loss, macrocephaly). For generation of cell lines, tumors from RCAS-injected mice were harvested, and mechanically and chemically homogenized. Single cell suspensions were cultured in NeuroCult Basal Medium containing NeuroCult Proliferation Supplement, 20 ng/ml EGF, 10 ng/ml basic FGF, 2 μg/ml heparin (Stemcell Technologies), 50 units/ml penicillin, and 50 μg/ml streptomycin (Thermo Fisher Scientific).

### Growth assay

Cell growth was monitored using Cell Titer 96 AQueous One proliferation assay (MTS) (Promega, Madison, WI) as per manufacturer’s instructions. Cells were plated onto 96-well plates as triplicates at the following densities: CT2A – 2000 cells/ well; M059J – 5000 cells/ well. Growth was measured at 24, 48, 72 and 96hrs after cell plating by combining cells in media with CellTiter 96 Aqueous Solution reagent, incubating at 37°C for 3hrs and measuring absorbance at 490nm. Wells containing media only were included for background subtraction.

### Cell treatments

Innate immune agonists obtained from Invivogen (San Diego, CA) include poly(I:C) LMW (Cat no: tlrl-picw) and poly(I:C) HMW (Cat no: tlrl-pic). BAY1436032 was obtained from MedChem Express (Monmouth Junction, NJ) and maintained as 10mM stocks in 100% ethanol. Cell lines treated with 10μg/ml poly(I:C) LMW or HMW for 4hrs or 24hrs were harvested for protein extraction at either 4hr or 24hr timepoints, while conditioned media was collected at the 24hr timepoint. Cells treated with 1μM BAY1436032 or with ethanol (vehicle control) every day for 3 days were harvested 24hours after the final dose for protein extraction, while conditioned media collected at the same timepoint was used for measuring D2HG and cytokine levels.

### Protein extraction and Western blotting

Monolayer cell cultures were washed in ice-cold 1X PBS twice and lysed in RIPA buffer containing Halt Protease and Phosphatase Inhibitor cocktail (Thermo Scientific) and Benzonase (Milliprore Sigma, Burlington, MA). Cell lysates were incubated on ice for 15 mins, followed by centrifugation at 14,000 rpm for 15 mins at 4°C. Supernatants were used to determine protein concentrations by Bradford assay. Twenty micrograms of protein were used in gel electrophoresis and western blotting. PVDF membranes were incubated with primary antibody overnight, followed by secondary antibody for 1 hour and processed for protein detection using SupersignalTM West Pico PLUS (Thermo Scientific) or Immobilon Western chemiluminescent substrate (Millipore Sigma). Images were captured and analyzed using ImageLab (Biorad, Hercules, CA). Primary and secondary antibodies used in this study are listed in Supplementary Table 2.

### Cytokine/chemokine analysis

Cytokine levels in conditioned media from cells were measured using Legendplex (Biolegend, San Diego, CA) as per the manufacturer’s instructions on a BD Fortessa X-20 flow cytometer. For CT2A and RCAS/Ntv-a samples, the mouse anti-virus response panel (Cat #: 740621) was used. For M059J samples, the human anti-virus response (Cat #: 740390) and pro-inflammatory chemokine panels (Cat #: 740984) were used. Samples were subjected to a 1:10 dilution to quantify levels of CCL5 and CXCL10 within the linear detection range. Legendplex Data Analysis software (https://legendplex.qognit.com) was used to calculate cytokine concentrations. Only cytokines with detectable induction upon treatment are shown.

### Flow cytometry analysis

Flow cytometry analysis on tumor bearing hemispheres harvested on day 14 post implant from C57BL/6 mice implanted with CT2A CRISPR control, *Atrx* KO-A or KO-B cells was performed as previously reported^13^. An additional step to remove myelin by overlaying on 20% Percoll (MP Biomedicals) solution in HBSS was included. Single cell suspensions stained with Zombie-Aqua (Biolegend, 1:500), were then stained with the following antibody panels (all Biolegend unless otherwise specified at 1:100 dilution): panel 1: CD45.2-BUV395 (BD Biosciences), CD3-FITC, CD19-FITC, NK1.1-BV421, CD11b-BV711, and Ly6G-PE; panel 2: CD45.2-BUV395, CD3-PE, CD4-FITC, CD8-BV421, PD1-PEcy7, and Tim3-BV711. Antibody-stained cell suspensions were analyzed using a BD Fortessa X-20 flow cytometer. Data were analyzed using Flow-Jo v10.8.1 (BD Biosciences). Gating strategies are included in Supplementary Fig. 6.

### D-2-hydroxyglutarate (D2HG) measurements

Cell pellets resuspended in PBS or conditioned media were used to quantify D2HG by liquid chromatography/ electrospray ionization tandem-mass spectrometry (LC-ESI-MS/MS) as published previously^42–44^, with modifications to accommodate equipment and sample matrix. D2HG concentrations in cell pellets were normalized to total protein.

### RNA extraction and gene expression analysis

RNA was extracted from monolayer cell cultures using miRNeasy Mini kit (Qiagen, Germantown, MD) according to manufacturer’s instructions. mRNA sequencing on CT2A *Atrx* KO lines and M059J *ATRX* KO lines was performed by Genewiz Azenta Life Sciences (South Plainfield, NJ) as per their workflow. cDNA libraries were sequenced using a HiSeq 2X platform (Illumina) to generate 150bp pair-end reads. mRNA sequencing on CT2A MSCV EV control or *Idh1*^R132H^ lines was performed by the Sequencing and Genomics Technologies Core Facility at Duke University. cDNA libraries generated using the Kapa stranded mRNA Kit (Roche Kapa Biosystems, Indianapolis, IN) were pooled to equimolar concentrations and sequenced on the NovaSeq 6000 S-Prime flow cell to produce 100 bp paired-end reads. Subsequent data processing, differential expression analysis, gene set enrichment analysis (GSEA) was performed as reported previously^45^.

To perform single sample gene set enrichment analysis (ssGSEA), log normalized TPM counts were input into GSVA v1.47.0 with method set to “ssgsea” and a list of custom gene sets. Enrichment scores were then Z-score transformed via the scale function in R, with the matrix transposed to scale across sample IDs. Heatmaps were generated via the package ComplexHeatmap v2.14.0.

For RCAS tumor sequencing, RNA extraction was performed with Qiagen RNeasy Plus kit, according to manufacturer’s instructions. Subsequent processing and sequencing occurred at the MD Anderson Advanced Technology and Genomics Core, (ATGC). Following Agilent BioAnalyzer quality assessment, libraries were generated using Truseq library preparation kits (Illumina) and samples run on the HiSeq4000 platform in a 76bp pair-end sequencing format. The raw fastq files were subjected to FASTQC analysis for quality control analysis, followed by alignment to mouse mm9 genome using RNASTAR (version 2.7.8a). Raw transcript counts were generated using HTseq-count tool (version 0.9.1), followed by Principal Component Analysis (PCA). Differentially expressed genes (DEGs) were calculated using limma-voom (version 3.44.3) and were later subjected to Gene Set Enrichment Analysis (GSEA) analysis using a desktop version of the analysis tool.

### TCGA data

Gene mutation data from the CNS/Brain TCGA LGG PanCancer Atlas Study from cBioportal was used to generate an Oncoprint output for a select number of immune genes. RSEM batch normalized counts (RNAseq) for 410 samples from the same dataset were stratified by *IDH* and *ATRX* mutation status, and 1p/19q codeletion status. Log normalized counts was subject to ssGSEA as described above for cell line RNAseq data.

### scRNA-Seq data extraction & analysis

scRNA-Seq public datasets *IDH*-mutant astrocytoma (GSE89567)^26^ and oligodendroglioma (GSE70630)^27^ were downloaded from NCBI GEO. Seurat’s v4.3.0 standard pre-processing workflow was used with clustering resolution determination aided by the R package clustree v0.5.0. Immune and tumor canonical marker genes were utilized along with differential expression lists to annotate each dataset’s clusters. Clusters annotated as immune cells or tumor cells were identified and integrated using Seurat’s integration workflow. Differential gene expression analysis was then performed via the FindAllMarkers function. Differential expression results were ranked by log2FC and then input into the GSEA R package fgsea v1.22.0 along with the Hallmark gene set collection.

### Statistical analysis

Results for all experiments are expressed as mean ± SEM. The number of biological or technical replicates and associated statistical tests are indicated in the corresponding figure legends. Data plotting and statistical analysis were performed using GraphPad Prism v9. p-values less than 0.05 are considered statistically significant and significant differences are indicated by asterisks (*) or hashtags (#). Individual data points are shown for all graphs.

## Data and materials availability

All data generated during this study are included in this published article and supplementary data files. Data are also available from the corresponding authors upon reasonable request. All transcriptional data have been deposited in Gene Expression Omnibus (GEO) under accession numbers: GSE228242 (CT2A *Atrx*-KO cell lines), GSE228181 (CT2A *Atrx*-KO/ *Idh1* mutant cell lines), GSE228243 (M059J *ATRX*-KO cell lines), GSE Pending (RCAS Atrx-KO cell lines), GSE Pending (RCAS Atrx-KO tumors). Source data file is available with this paper.

## Acknowledgments

We thank Feng Zhang for providing the CRISPR-Cas9 lentiCRISPR v2 vector (Broad Institute, Boston, MA). We thank Duke’s Sequencing and Genomic Technologies Shared Resource for providing the mRNA-sequencing service, Duke’s Brain Tumor Center Biorepository and Database, Substrate services core research support and Duke’s Pathology Research Histology and Immunohistochemistry Laboratory for IHC staining.

We thank the MD Anderson Genomics Core Facility for mRNA-sequencing on the RCAS model. This work was supported by NIH R01 CA255788 (D.M.A, J.T.H and M.C), R01 CA240338 (J.T.H.) and American Cancer Society’s Research Scholar’s Grant, RSG-16-179-01-DMC (J.T.H.). Research reported in this publication was supported by the National Center for Advancing Translational Sciences of the National Institutes of Health under Award Numbers TL1TR003169 and UL1TR003167 (B.T.W) and NCI Cancer Center Support Grant (CCSG), P30CA014236 (I.S.). The content is solely the responsibility of the authors and does not necessarily represent the official views of the National Institutes of Health. We would like to acknowledge V Foundation for Cancer Research, Jewish Communal Fund Grant and Brain Tumor Research Charity Grant for funding provided to D.M.A. and the Ben and Catherine Ivy Foundation and the Brockman Foundation for funding provided to J.T.H.

## Author contributions

Conceptualization and design: S.H., B.T.W., M.S.W., M.C.B., M.L.B., D.M.I., S.D., S.T.K., M.G.C., J.T.H., D.M.A.

Development of methodology: S.H., B.T.W., M.S.W., M.C.B., M.L.B., D.M.I., S.D., S.T.K., Y.H., J.T.H., D.M.A.

Acquisition of data: S.H., B.T.W., M.C.B., M.L.B., D.M.I., K.R., R.F., J.H., S.D., E.A.G., A.B., A.M., C.F., M.S., M.R., I.S., S.T.K., Y.H.

Analysis and interpretation of data: S.H., B.T.W., C.J.P., M.S.W., M.C.B., M.L.B., D.M.I., S.D.,

K.S., G.Z., P.B.M., H.R., S.T.K., Y.H., M.G.C., J.T.H., D.M.A.

Writing, review and/or revision of manuscript: S.H., B.T.W., C.J.P., M.S.W., M.C.B., M.L.B., M.G.C., J.T.H., D.M.A.

Administrative, technical, or material support: S.H., B.T.W., M.L.B., J.T.H., D.M.A. Study supervision: J.T.H., D.M.A.

## Competing interests

The authors declare no competing interests.

Materials & correspondence:

Correspondence and material requests should be addressed to D.M.A. and J.T.H.

