## Supplementary data for "Interplay between *ATRX*and *IDH1* mutations governs innate immune responses in diffuse gliomas"

### Supplementary figure 1

**a** Innate immune gene sets

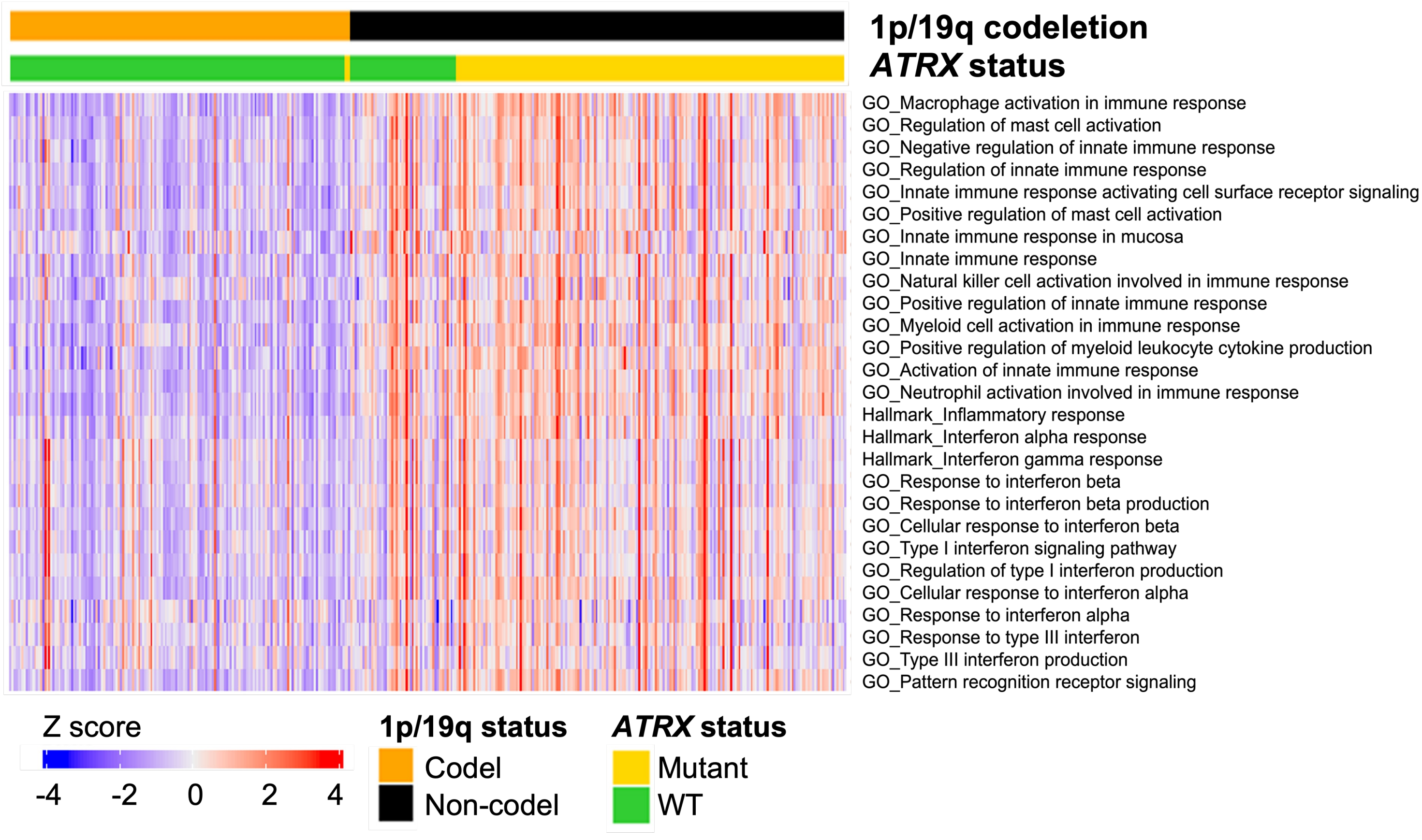


**b** General & adaptive immune gene sets

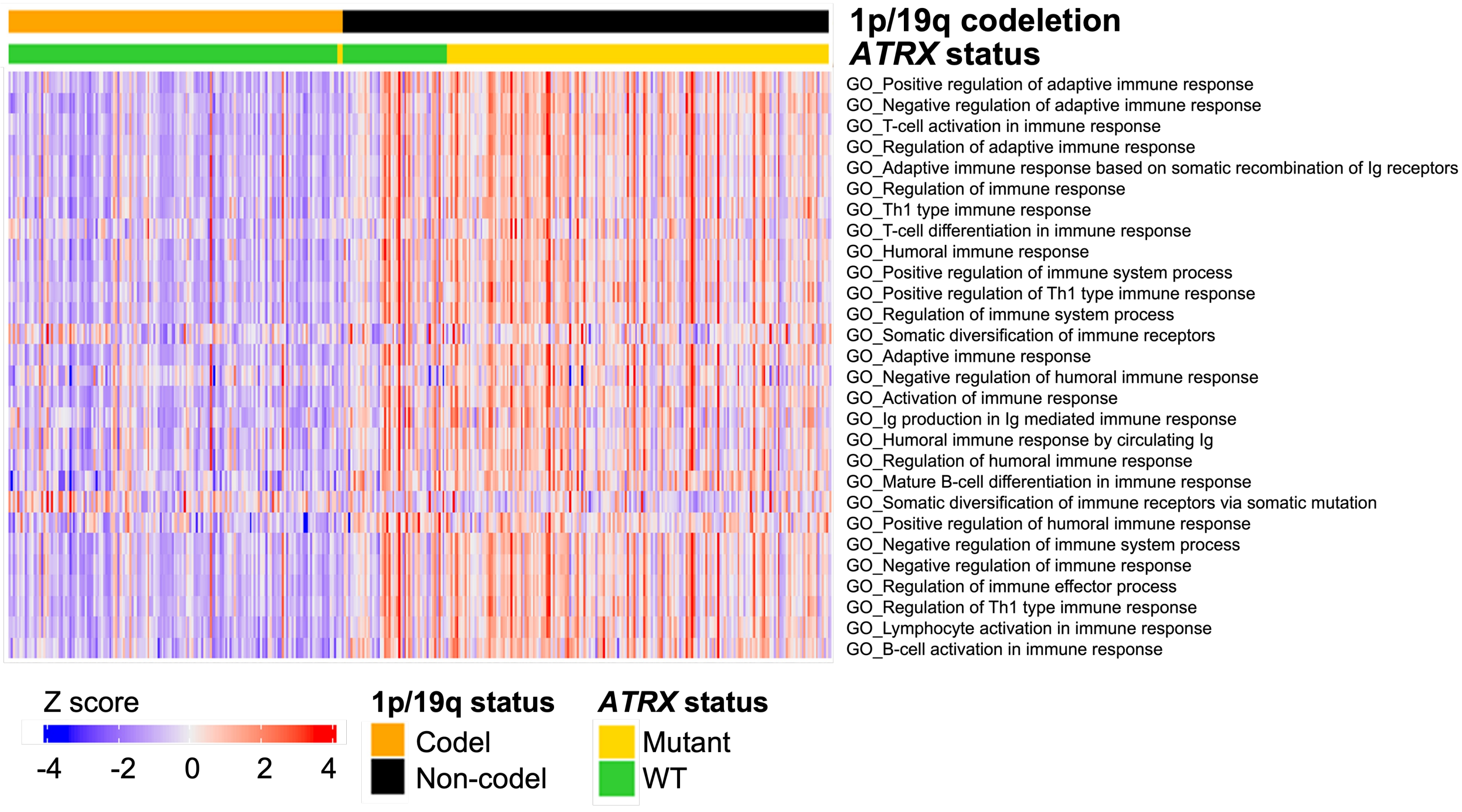

**Supplementary figure 1 – supporting figure 1a. Astrocytomas are immunologically engaged compared to oligodendrogliomas.**

(**a, b**) Heatmap from ssGSEA showing enrichment for various innate immune-related (**a**) and general and adaptive immune-related (**b**) gene sets in 1p-19q noncodel/ *IDH* mutant/ *ATRX*-mutant astrocytomas (n=191) compared to 1p-19q codel/ *IDH* mutant/ *ATRX*-WT oligodendrogliomas (n=164). A relatively smaller number of 1p-19q codel/ *IDH* mutant/ *ATRX*-mutant (n=3) and 1p-19q noncodel/ *IDH* mutant/ *ATRX* WT (n=52) are also included in the analysis for comparison.

### Supplementary figure 2

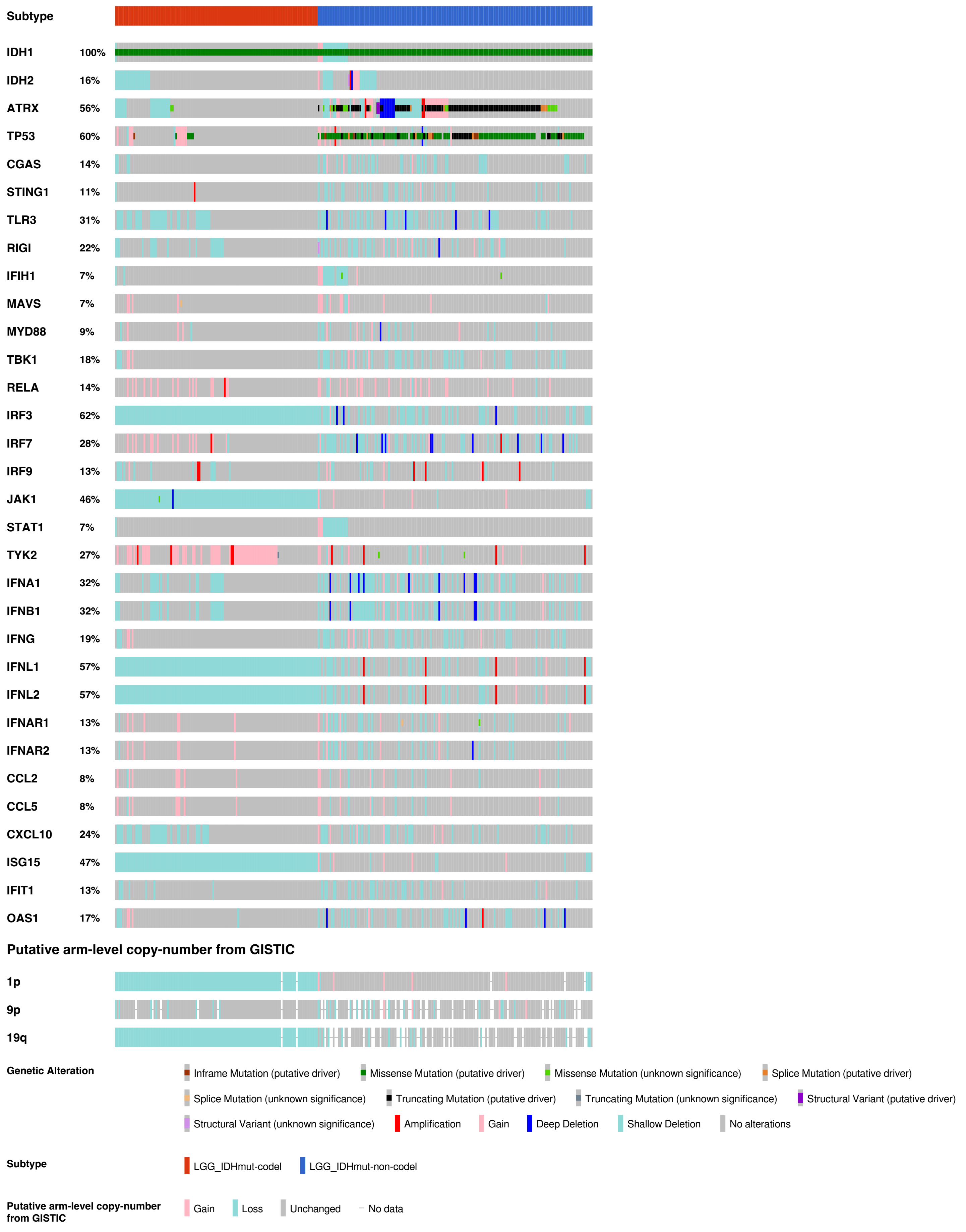

**Supplementary figure 2 – supporting figures 1a and b. *IDH*-mut astrocytomas exhibit lower frequency of genetic alterations in immune-related genes compared to *IDH*-mut oligodendrogliomas.** Oncoprint output from cBioportal analyzing *IDH* mutant-LGGs for mutation status of indicated genes. Datasets from *IDH*-mutant 1p/19q codel subtype are compared with *IDH*-mutant 1p/19q noncodel subtype.

### Supplementary figure 3

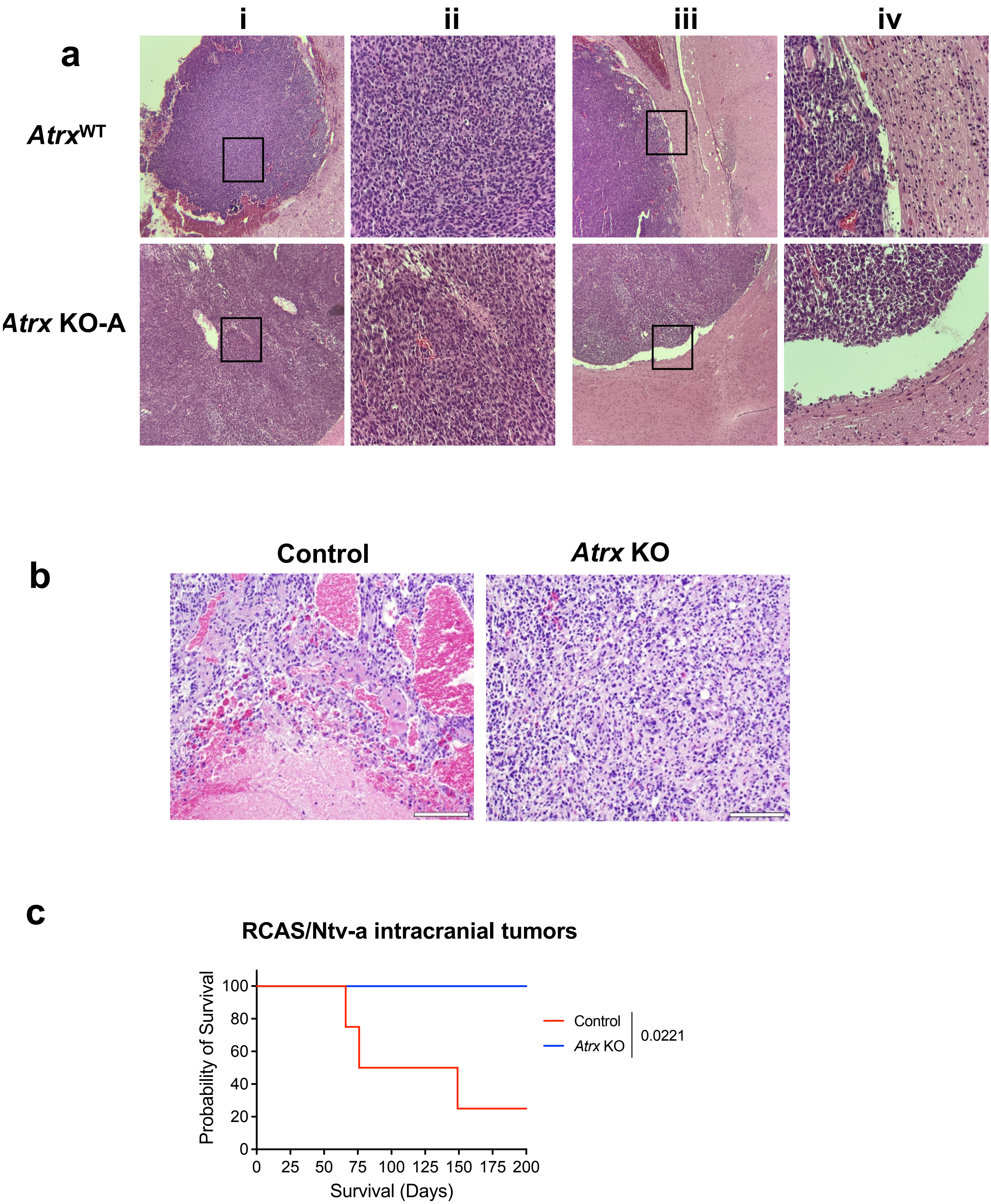

**Supplementary figure 3 – supporting figures 2a and b. Histology from *Atrx*$^{WT}$ and *Atrx*-KO CT2A and RCAS/Ntv-a tumors.**

(**a**) H&E staining performed on end-stage CT2A CRISPR control (*Atrx* WT), *Atrx* KO-A or KO-B tumors. (**I, iii**) represent images at 4X; (**ii, iv**) represent images at 20X. (**b**) H&E staining performed on end-stage *Atrx*$^{fl/fl}$ (*Atrx* KO) or *Atrx*$^{+/+}$ (Control) tumors. **(c)** Kaplan Meir survival curves for C57BL/6 mice bearing intracranial RCAS/ Ntv-a *Atrx*$^{fl/fl}$ (*Atrx* KO) (n=5) and *Atrx*$^{+/+}$ (Ctrl) (n=4) tumors generated by re-injecting *Atrx*$^{+/+}$ (Ctrl) or *Atrx*$^{fl/fl}$ (*Atrx* KO) cell lines derived from the de-novo model. P-values representing group comparisons were calculated using log-rank test.

### Supplementary figure 4

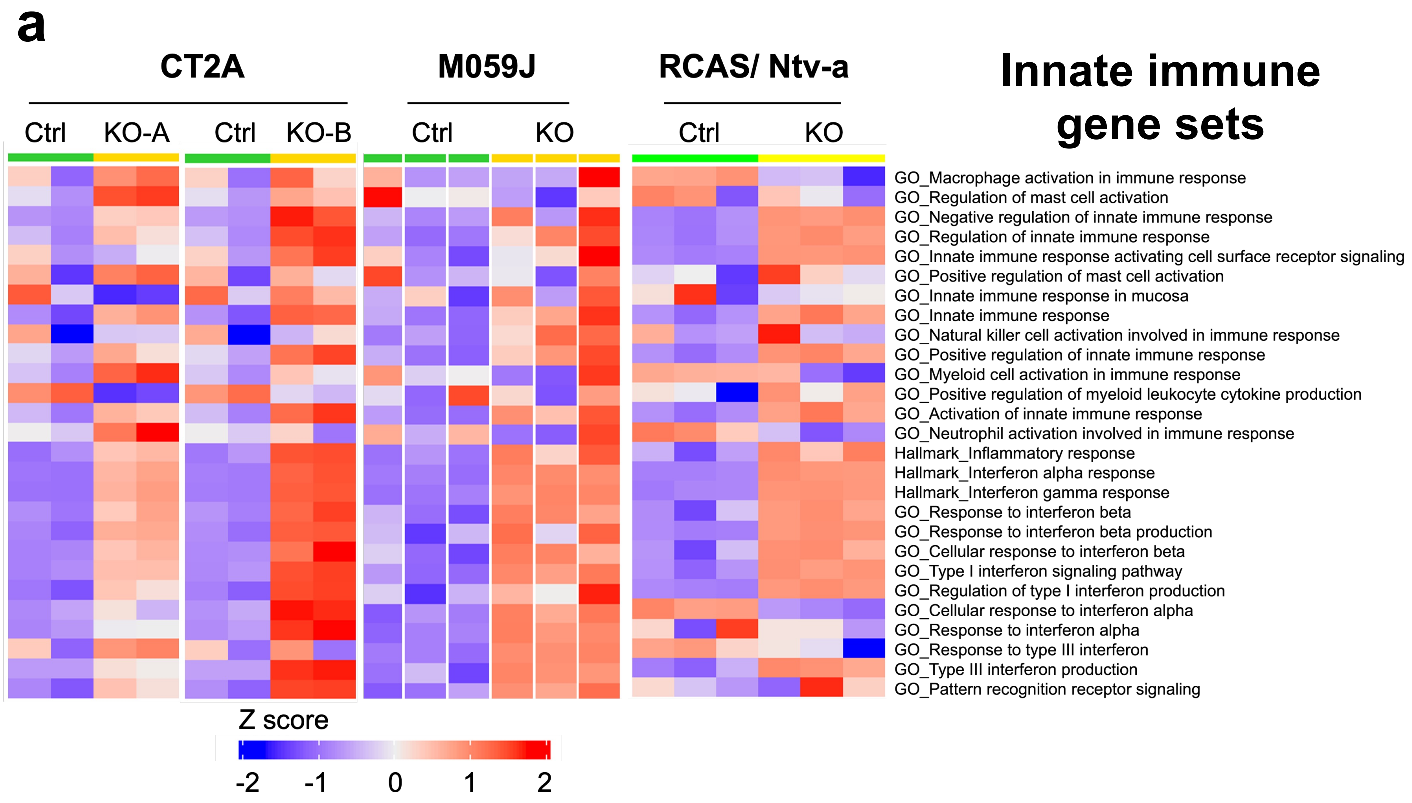


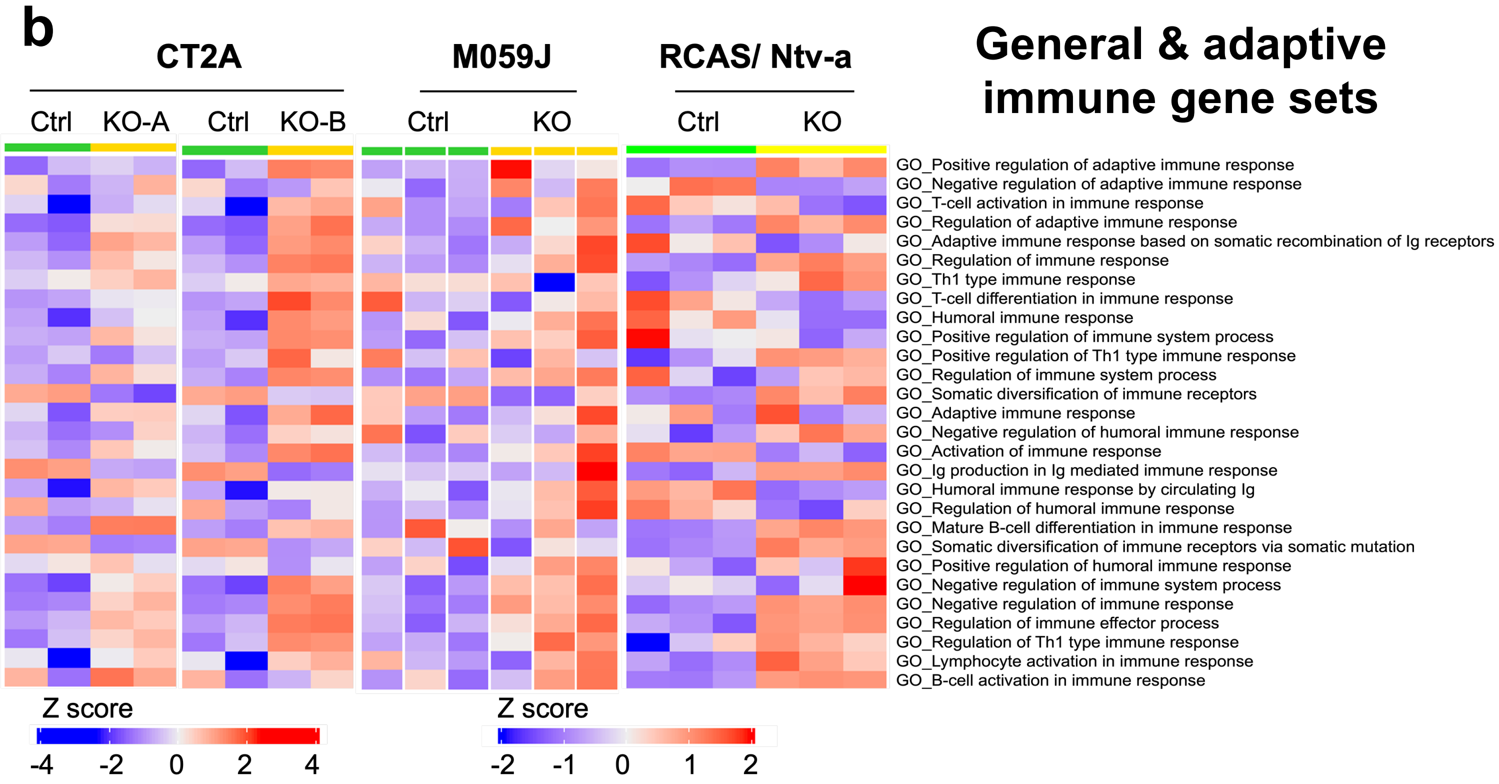

**Supplementary figure 4 – supporting figure 2d. ATRX loss is associated with a pro-inflammatory signaling.**
(**a, b**) Heatmap from ssGSEA showing enrichment for various innate immune-related (**a**) and general and adaptive immune-related (**b**) gene sets in CT2A *Atrx* KO-A and KO-B clones, M059J *ATRX* KO cells and RCAS/Ntv-a *Atrx* KO compared to their respective *ATRX*-WT counterparts. n=2 technical replicates per cell line for CT2A expression data; n= 3 technical replicates per cell line for M059J expression data; n=3 biological replicates (3 consecutive cell passages) per cell line for RCAS/Ntv-a cell line expression data.

### Supplementary figure 5

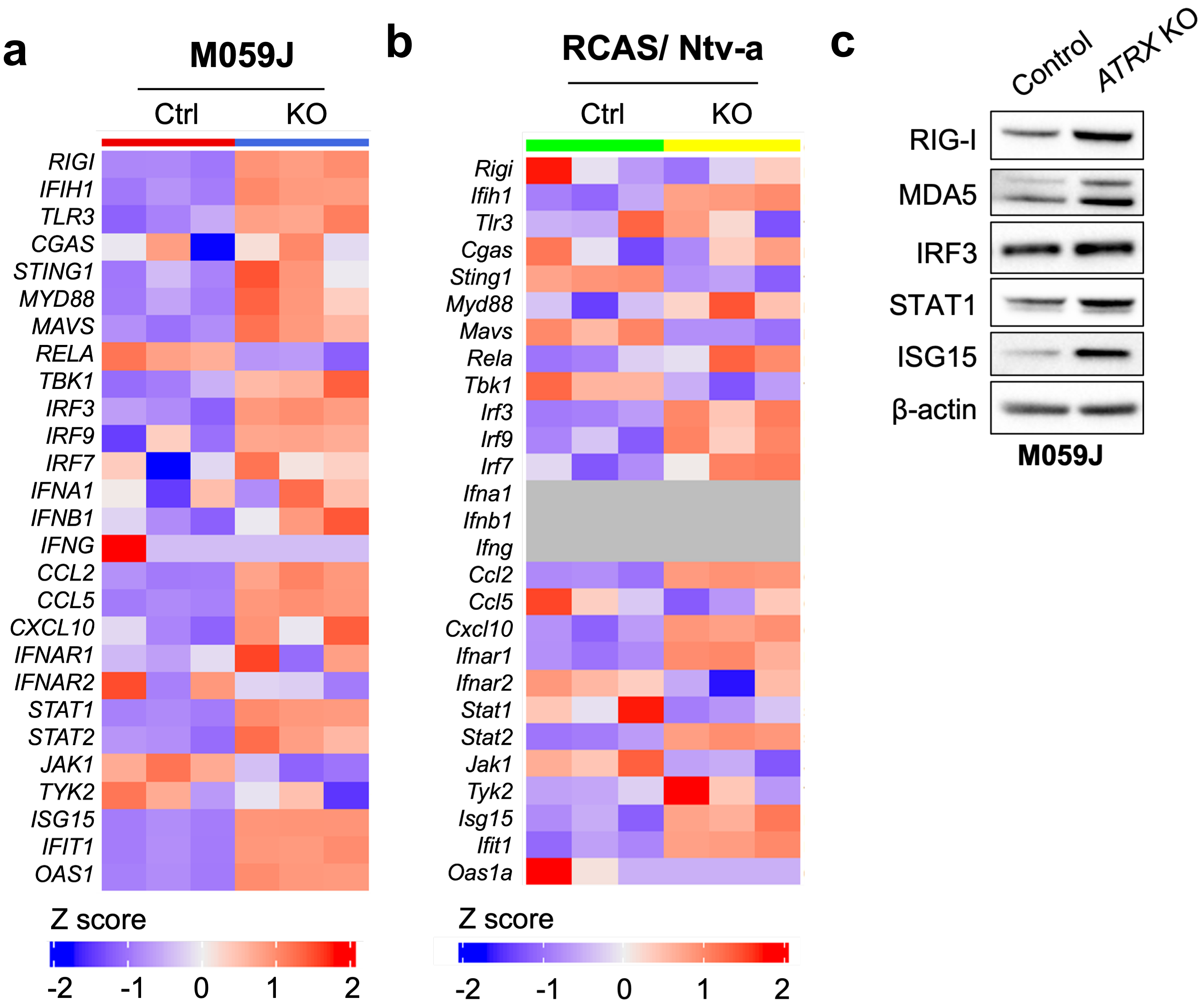


**Supplementary figure 5 – supporting figures 3a and b. ATRX loss is associated with increased baseline gene and protein expression*.***

(**a, b)** Heatmap showing differential expression of immune-related genes in M059J CRISPR control (Ctrl) and *ATRX* KO cells (**a**) and RCAS/Ntv-a *Atrx*$^{+/+}$ (Ctrl) and *Atrx*$^{-/-}$ (KO) cells (**b**). n= 3 technical replicates per cell line for M059J expression data; n=3 biological replicates (3 consecutive cell passages) per cell line for RCAS/Ntv-a cell line expression data.

(**c**) Western blot using lysates from M059J CRISPR ctrl (Ctrl) and ATRX KO cells (KO) screened for proteins involved in innate immune signaling. β-actin serves as the loading control.

### Supplementary figure 6

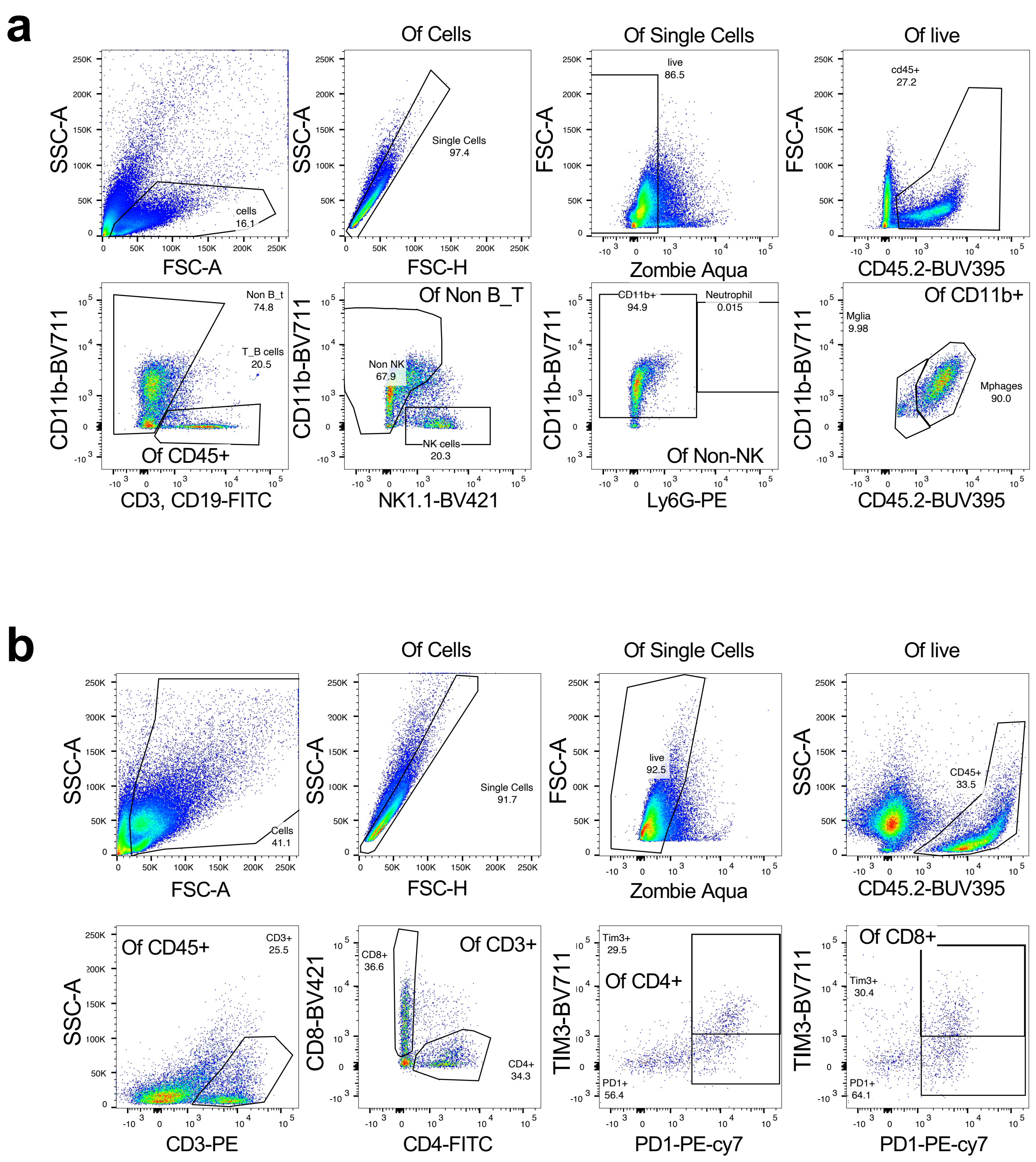


**Supplementary figure 6 – supporting figure 4a. Representative gating strategy for analysis of *Atrx*-KO tumors.**
Gating strategy for analysis of tumor associated myeloid (**a**) and T cells (**b**) along with relevant phenotypes by flow cytometry.

### Supplementary figure 7

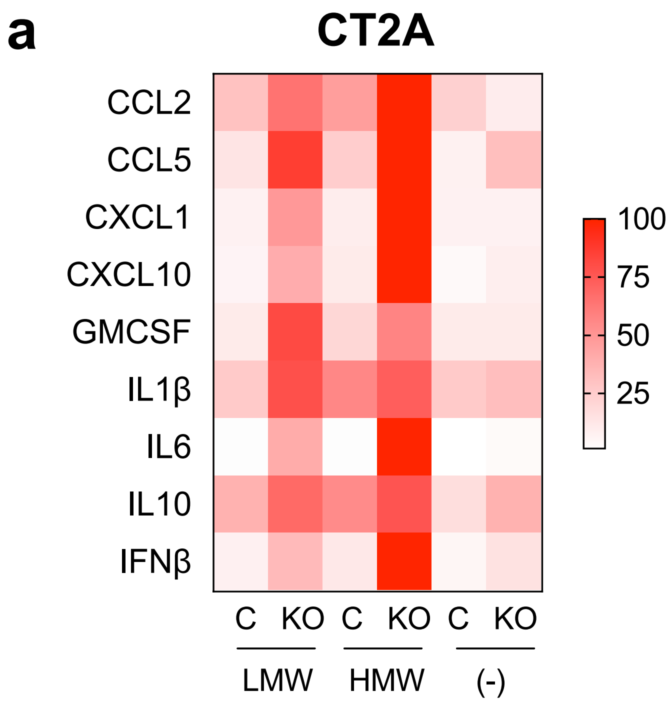


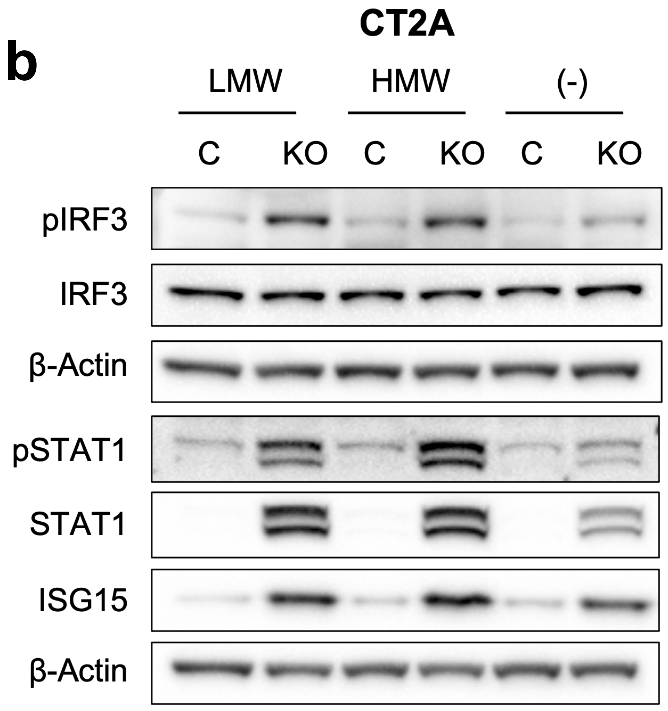


**Supplementary figure 7 – supporting figures 5a and d. ATRX depletion sensitizes cells to poly(I:C), a dsRNA agonist.**
(**a**) Cytokine levels in conditioned media from CT2A CRISPR control and *Atrx* KO-B cells, treated with poly(I:C) LMW or HMW for 24hrs. Supernatant cytokines were analyzed by cytokine bead arrays for antiviral and proinflammatory cytokines. For heatmap generation, maximum values for each cytokine were set to 100%. Only cytokines and signaling proteins with observed induction after treatment with poly(I:C) are included. n=2 biological replicates. (**b**) Western blot using lysates from CT2A CRISPR control and *Atrx* KO-B cells treated with poly(I:C) LMW or HMW for 4hrs or 24hrs, screened for pIRF3/IRF3, pSTAT1/STAT1 and ISG15 involved in innate immune signaling. β-actin serves as the loading control.

### Supplementary figure 8

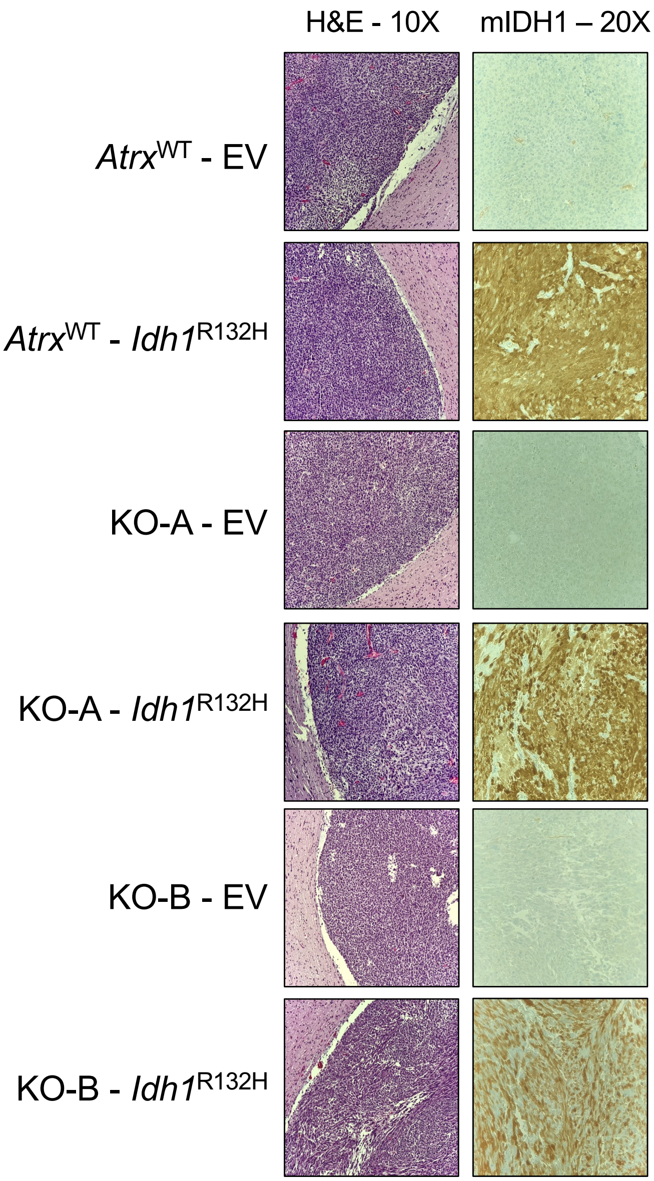


**Supplementary figure 8 – supporting figure 6d.** H&E and IDH1$^{R132H}$ staining of CT2A *Idh1*$^{WT}$ or *Idh1*$^{R132H}$ tumors harvested when mice were moribund.

### Supplementary figure 9

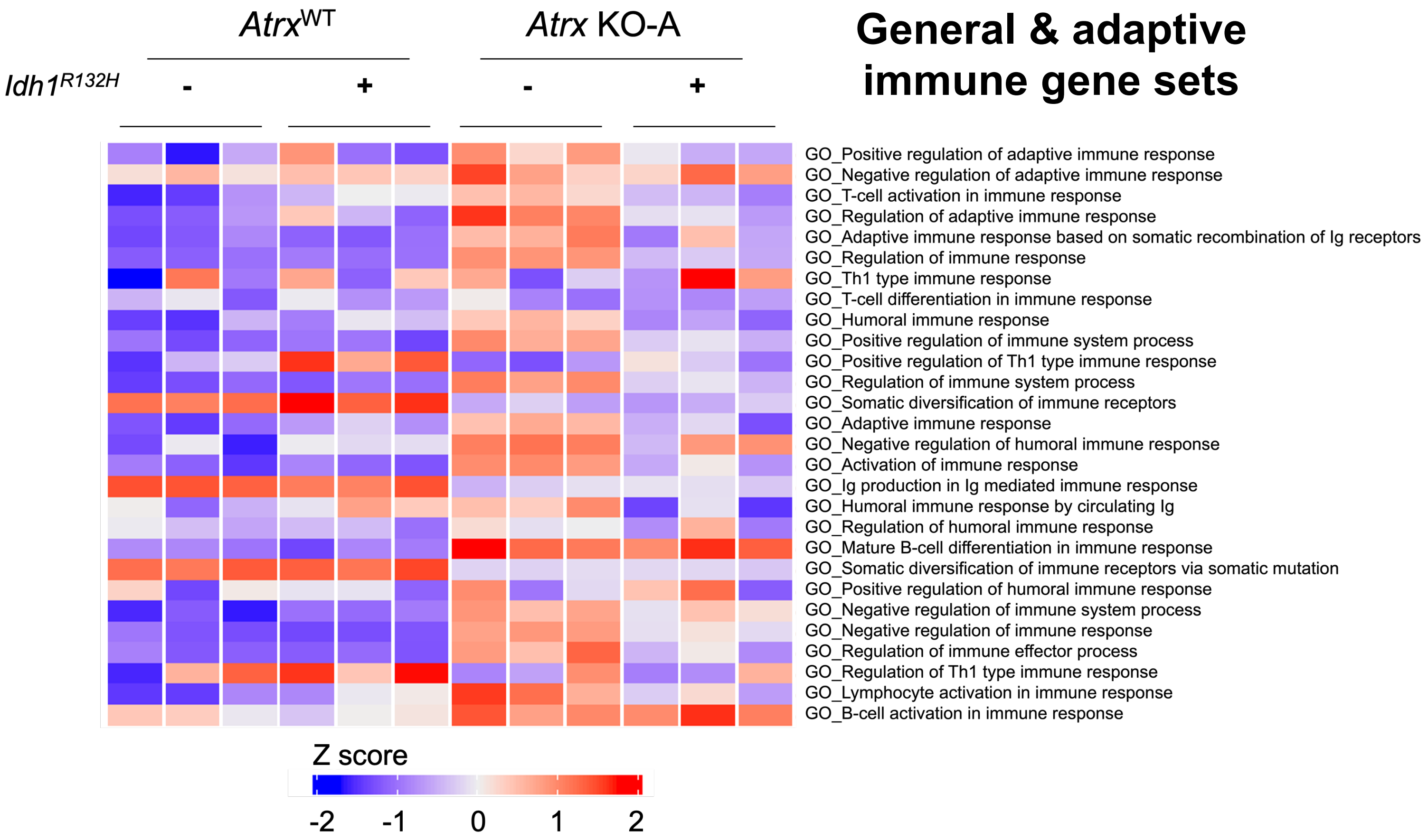


**Supplementary figure 9 – supporting figure 7a. Idh1$^{R132H}$ co-expression in Atrx-deficient KO-A cells dampens baseline innate gene expression.**

Heatmap from ssGSEA showing loss of enrichment of various general and adaptive immune-related GO terms in *Atrx* KO-A/ *Idh1$^{R132H}$* cells compared to *Atrx KO*-A/ *Idh1$^{WT}$* cells. RNA was isolated from cells cultured for 72hrs. n=3 technical replicates per cell line.

### Supplementary figure 10

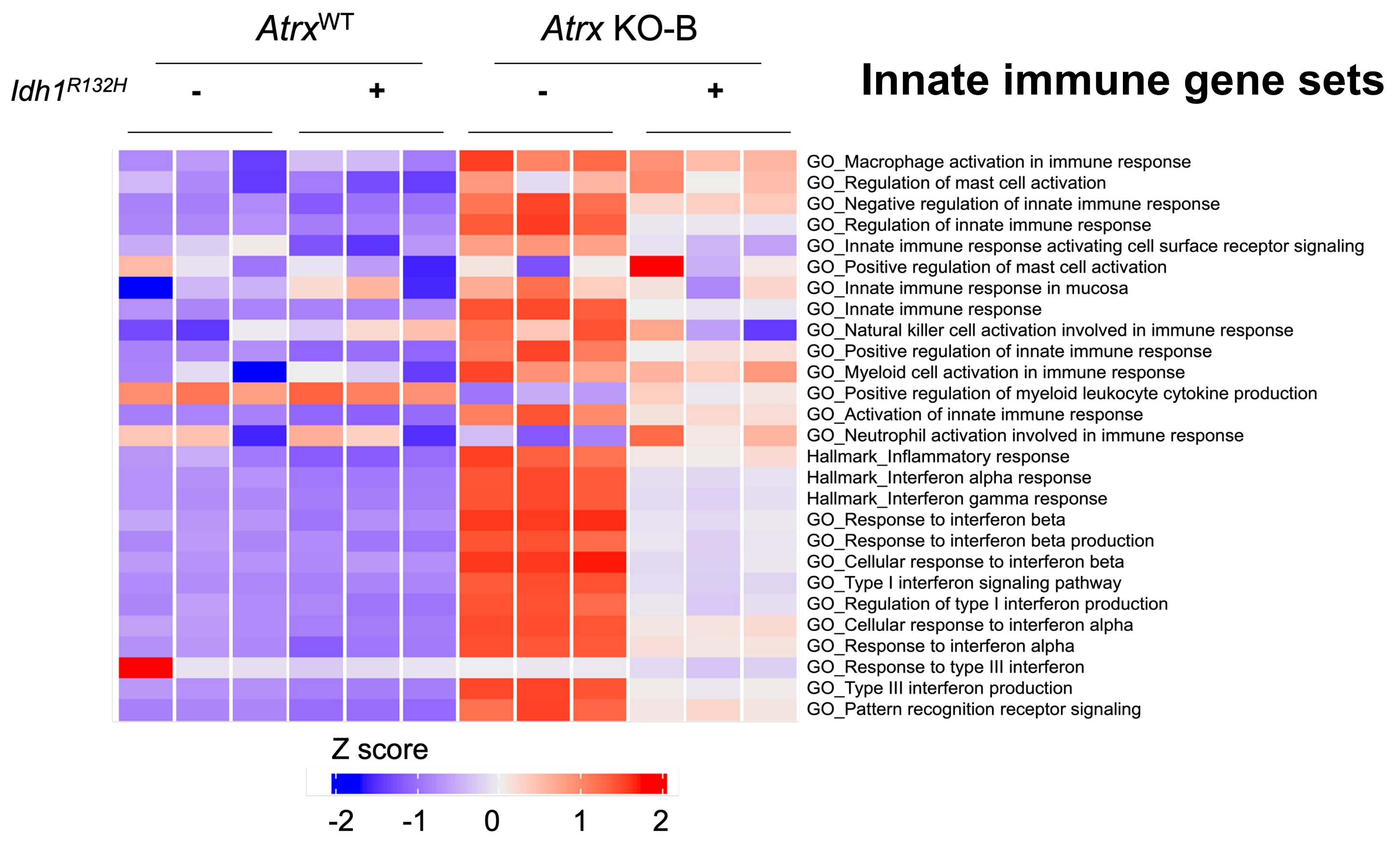


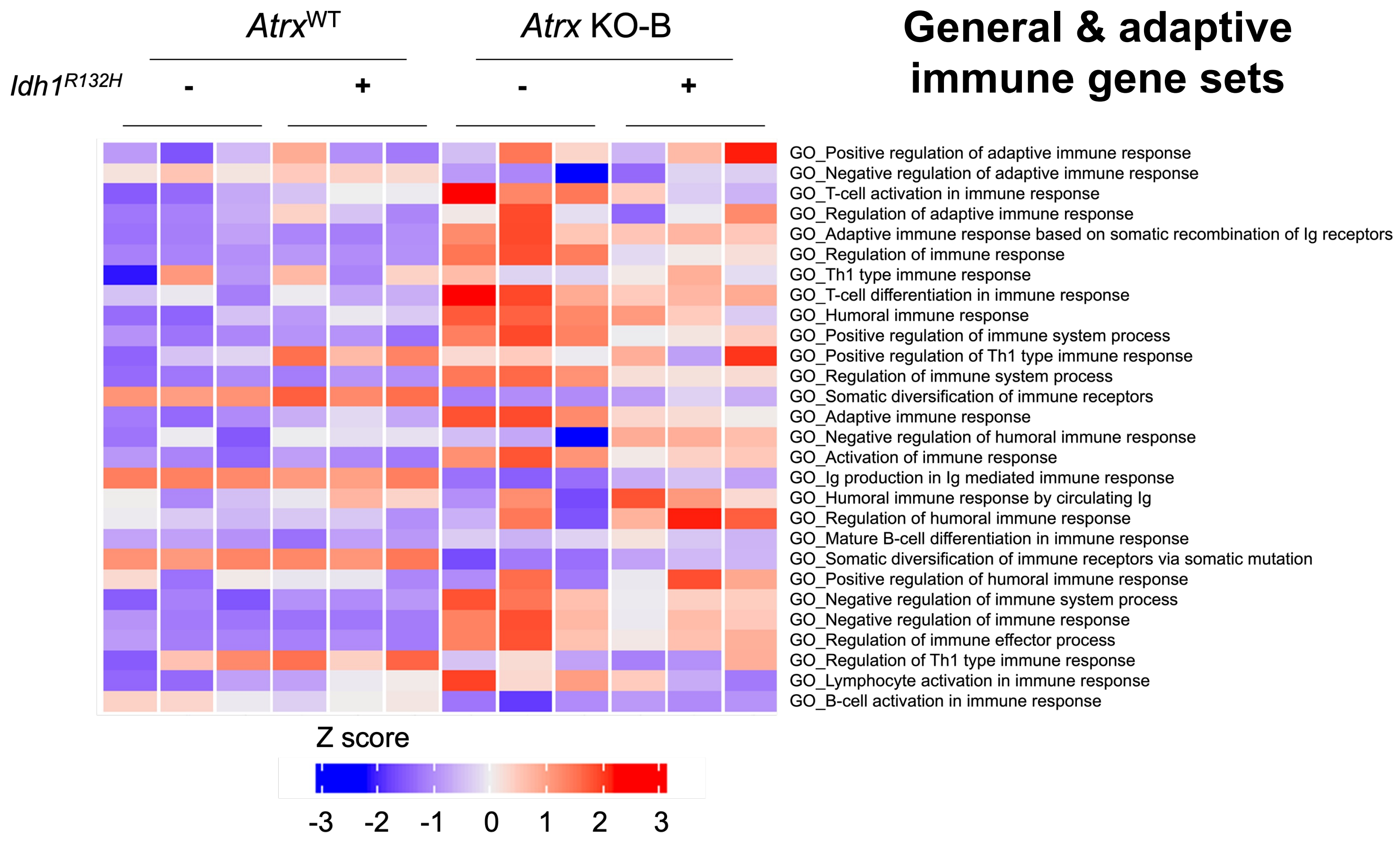

**Supplementary figure 10 – supporting figure 7a. Idh1$^{R132H}$ co-expression in Atrx-deficient KO-B cells dampens baseline innate gene expression**.

Heatmap from ssGSEA showing loss of enrichment of various innate and adaptive immune-related GO terms in *Atrx* KO-B/ *Idh1$^{R132H}$* cells compared to *Atrx KO*-B/ *Idh1$^{WT}$* cells. RNA was isolated from cells cultured for 72hrs. n=3 technical replicates per cell line.

### Supplementary figure 11

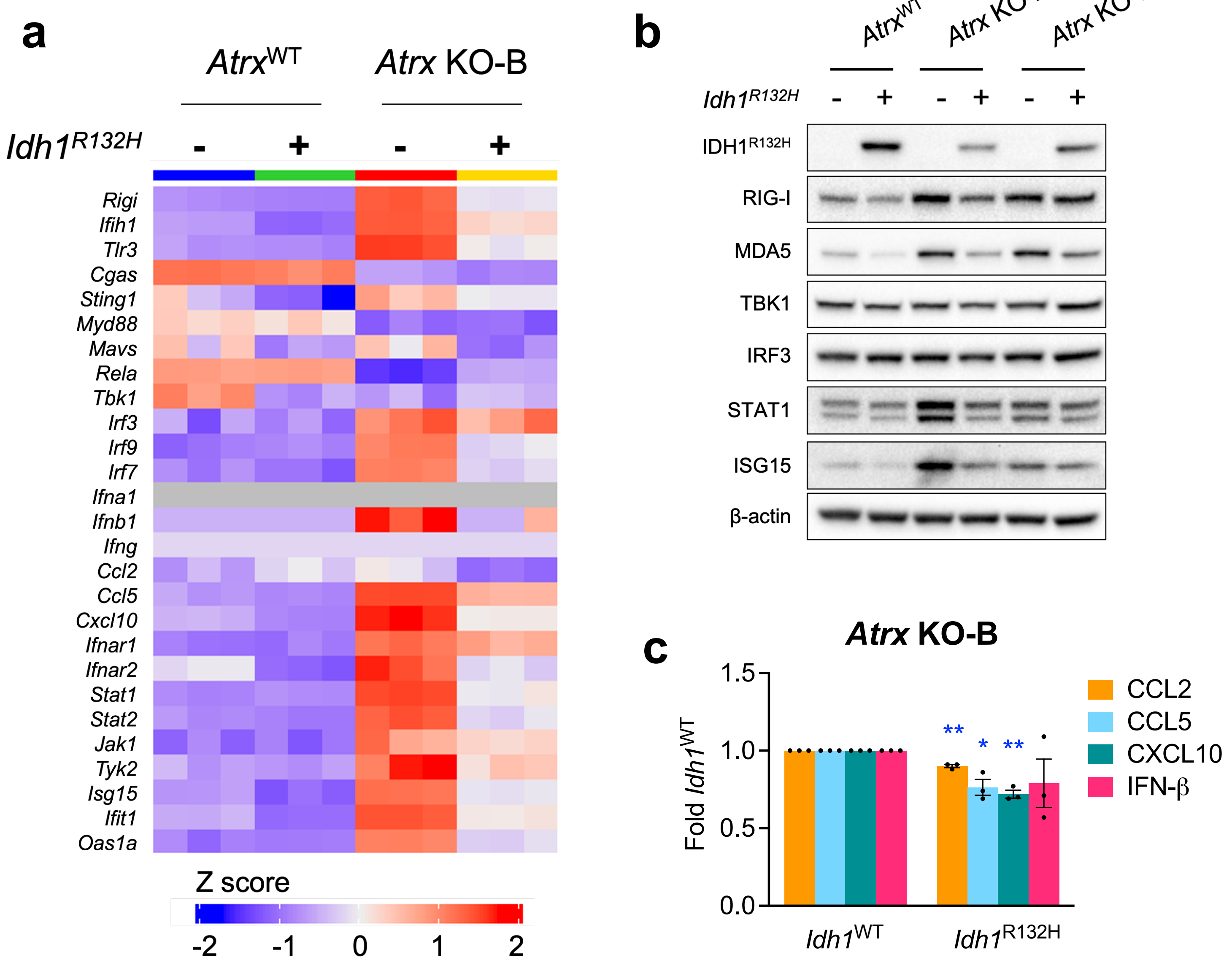

**Supplementary figure 11 – supporting figures 7b-7d. Idh1$^{R132H}$ co-expression in Atrx-deficient cells dampens baseline innate protein expression and cytokine secretion**.

(**a**) Heatmap showing differential expression of immune-related genes in CT2A *Atrx*$^{WT}$ and *Atrx* KO-B cells with or without *Idh1*$^{R132H}$. n=3 technical replicates per cell line. (**b**) Lysates from CT2A CRISPR control (*Atrx*$^{WT}$), *Atrx* KO-A and *Atrx* KO-B cells expressing exogenous Idh1$^{R132H,}$ or empty vector cultured for 72hrs, were screened by Western blotting for various innate immune proteins. β -actin serves as the loading control. (**c**) Conditioned media from CT2A *Atrx* KO-B cells expressing exogenous Idh1$^{R132H}$ or empty vector cultured for 72hrs was assayed for antiviral and proinflammatory cytokines using a Legendplex assay kit. Data indicates fold change values normalized to corresponding *Idh1*$^{WT}$ sample, shown as mean $\pm$ SEM. n=3 biological replicates. Asterisks denote significant two-tailed t-tests. (*: $p<0.05$; **: $p<0.01$; ***: $p<0.001$).

### Supplementary figure 12

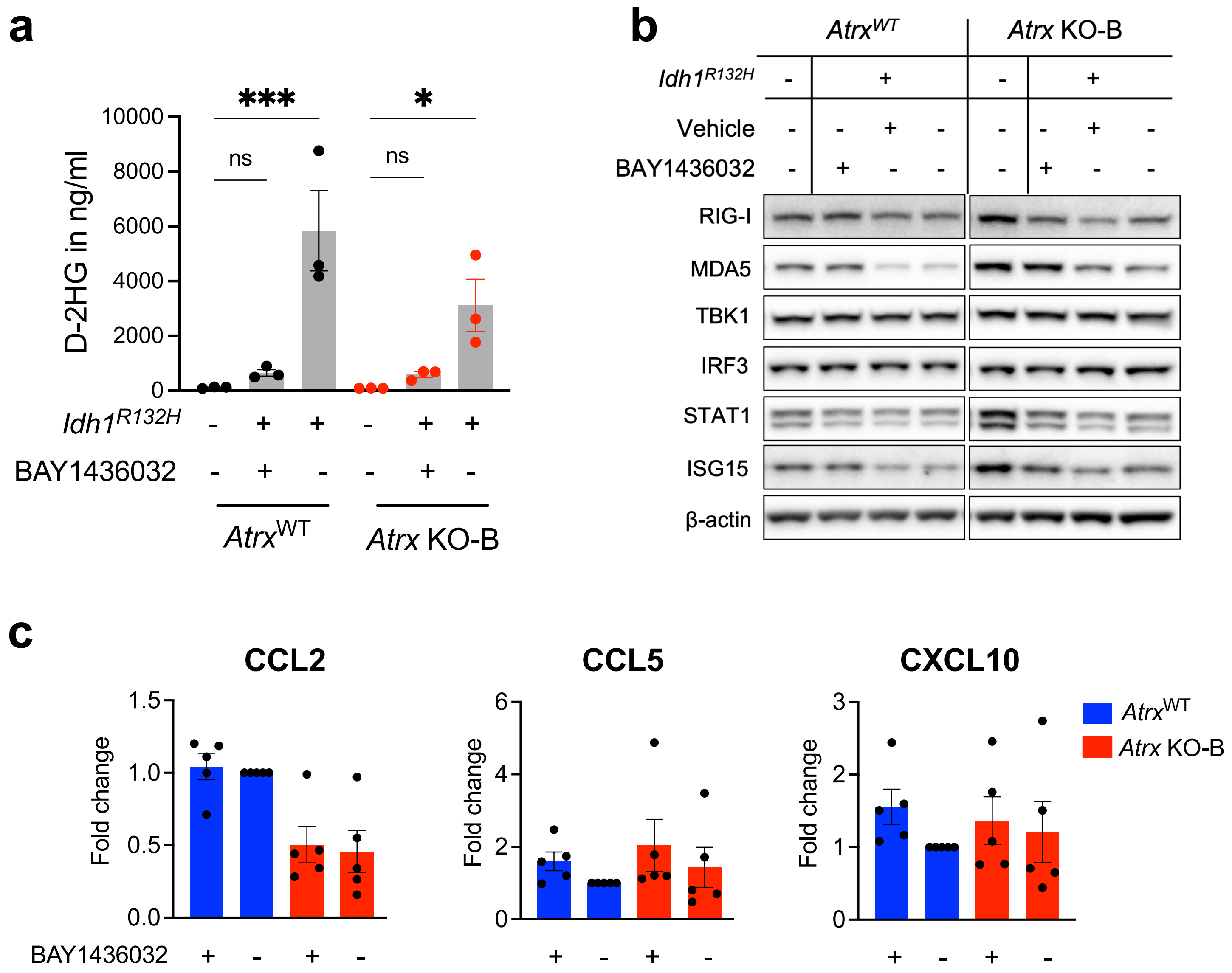

**Supplementary figure 12 – supporting figure 8. BAY1436032 partially reverses *IDH1$^{R132H}$*-mediated immunosuppression.**
(**a**) D-2HG levels in conditioned media from CT2A CRISPR control (*Atrx*$^{WT}$) or *Atrx* KO-B cells expressing IDH1$^{R132H}$, or empty vector treated with 1μM BAY1436032 or vehicle every day for 3 days, normalized to total protein. n=3 biological replicates. Asterisks indicate significant p-values from one-way ANOVA with Sidak post-hoc test. (*: $p<0.05$, **: $p<0.01$ ***: $p<0.001$) (**b**) Western blot using lysates from CT2A CRISPR control (*Atrx*$^{WT}$) or *Atrx* KO-B cells expressing Idh1$^{R132H}$ or empty vector that were treated with 1uM BAY1436032 or vehicle every day for 3 days, screened for proteins involved in innate immune signaling. β-actin serves as the loading control. (**c**) Cytokine/chemokine levels in conditioned media from CT2A CRISPR control (*Atrx*$^{WT}$) and *Atrx* KO-B cells expressing Idh1$^{R132H}$ treated with 1uM BAY1436032 or vehicle every day for 3 days. Supernatant cytokines were analyzed by cytokine bead arrays for antiviral and proinflammatory cytokines. Fold change values normalized to vehicle treated *Atrx*$^{WT}$ sample are plotted. N=3 biological replicates. One-way ANOVA with Tukey's post-hoc test did not reveal any significant differences between groups.

### Supplementary figure 13

**a**

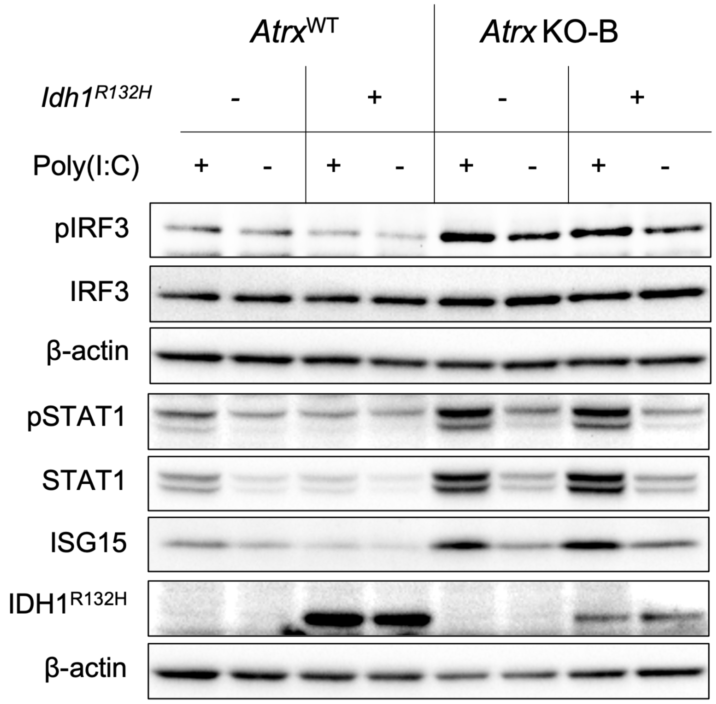


**b**

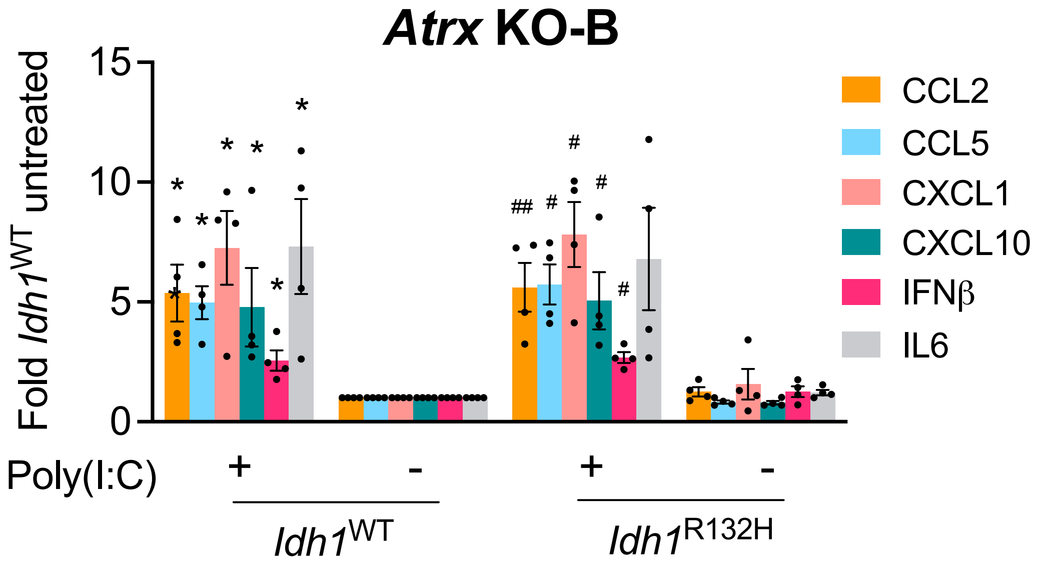

**Supplementary figure 13 - supporting main figure 9. *Atrx* KO/ *Idh1$^{R132H}$* cells retain sensitivity to poly(I:C).**

(**a**) Western blot using lysates from CT2A CRISPR control (*Atrx*$^{WT}$) or *Atrx* KO-B cells expressing Idh1$^{R132H}$ or empty vector that were treated with 10µg/ml poly(I:C) HMW for 4hrs or 24hrs and screened for proteins involved in innate immune signaling. (**b)** Cytokine levels in conditioned media from CT2A *Atrx* KO-B cells expressing Idh1$^{R132H}$ or empty vector treated with poly(I:C) HMW for 24hrs. Supernatant cytokines were analyzed by cytokine bead arrays for antiviral and proinflammatory cytokines. Fold change values normalized to untreated *Atrx* KO-B/ *Idh1*$^{WT}$ sample are shown as mean $\pm$ SEM. n= 4 biological replicates. Asterisks indicate significant p-values from one-way ANOVA with Tukey’s post hoc test.
