## Supplementary tables for "Interplay between *ATRX*and *IDH1* mutations governs innate immune responses in diffuse gliomas"

**Supplementary table 1:** Sequences of sgRNAs targeting human and mouse ATRX.

| | |
|---|---|
| sgRNA-hATRX-e9-sense | CACCGTGTTGGCAGGTTCATATTG |
| sgRNA-hATRX-e9-antisense | AAACCAATATGAACCTGCCAACAC |

| | |
|---|---|
| sgRNA-mATRX-ex9-1-sense | CACCGTGTAAAAACTACACCGTTG |
| sgRNA-mATRX-ex9-1-antisense | AAACCAACGGTGTAGTTTTTACAC |
| sgRNA-mATRX-ex9-2-sense | CACCGTATCTGACGATGAACACTC |
| sgRNA-mATRX-ex9-2-sense | AAACGAGTGTTCATCGTCAGATAC |

**Supplementary table 2**: Antibodies used in this study.

| Antibody | Company | Catalog number |
|---|---|---|
| ATRX (Human specific) | Cell signaling | 14820 |
| ATRX | Novus Biologicals | NBP1-32851 |
| ATRX | Cell signaling | 10321 |
| $IDH1^{R132H}$ | Dianova | DIA-H09 |
| $IDH1^{WT}$ | Cell signaling | 8137 |
| Phospho IRF3 (Ser396) | Cell signaling | 4947 |
| Phospho IRF3 (Ser396) | Thermo scientific | MA5-14947 |
| Total IRF3 (Human specific) | Cell signaling | 10949 |
| Total IRF3 | Biolegend | 655702 |
| Phospho STAT1 (Tyr701) | Cell signaling | 9167 |
| Total STAT1 | Cell signaling | 9172 |
| Total STAT1 | Cell signaling | 14994 |
| Total TBK1 | Cell signaling | 3504 |
| ISG15 | Cell signaling | 2743 |
| RIG-I | Cell signaling | 3743 |
| MDA5 | Cell signaling | 5321 |
| Vinculin | Sigma | V9264 |
| β-actin | Cell signaling | 4967 |
| β-actin, HRP conjugated | Cell signaling | 5125 |
| β-tubulin, HRP conjugated | Cell signaling | 5346 |
| Anti-rabbit IgG, HRP conjugated antibody | Cell signaling | 7074 |
| Anti-mouse IgG, HRP conjugated antibody | Cell signaling | 7076 |
| CD45.2-BUV395 | BD Biosciences | 564616 |
| CD3-FITC | BioLegend | 100204 |
| CD19-FITC | BioLegend | 152404 |
| NK1.1-BV421 | BioLegend | 108732 |
| CD11b-BV711 | BioLegend | 101242 |

| | | |
|---|---|---|
| Ly6G-PE | BioLegend | 127608 |
| CD3-PE | BioLegend | 100205 |
| CD4-FITC | BioLegend | 100510 |
| CD8-BV421 | BioLegend | 100738 |
